# Bond type and discretization of non-muscle myosin II are critical for simulated contractile dynamics

**DOI:** 10.1101/669382

**Authors:** D.B Cortes, M. Gordon, F. Nédélec, A.S. Maddox

**Affiliations:** UNC Chapel Hill; University of Cambridge, Sainsbury Laboratory

## Abstract

Molecular motors drive cytoskeletal rearrangements to change cell shape. Myosins are the motors that move, crosslink, and modify the actin cytoskeleton. The primary force generator in contractile actomyosin networks is non-muscle myosin II (NMMII), a molecular motor that assembles into ensembles that bind, slide, and crosslink actin filaments (F-actin). The multivalence of NMMII ensembles and their multiple roles have confounded the resolution of crucial questions including how the number of NMMII subunits affects dynamics, and what affects the relative contribution of ensembles’ crosslinking versus motoring activities. Since biophysical measurements of ensembles are sparse, modeling of actomyosin networks has aided in discovering the complex behaviors of NMMII ensembles. Myosin ensembles have been modeled via several strategies with variable discretization/coarse-graining and unbinding dynamics, and while general assumptions that simplify motor ensembles result in global contractile behaviors, it remains unclear which strategies most accurately depict cellular activity. Here, we used an agent-based platform, Cytosim, to implement several models of NMMII ensembles. Comparing the effects of bond type, we found that ensembles of catch-slip and catch motors were the best force generators and binders of filaments. Slip motor ensembles were capable of generating force but unbound frequently, resulting in slower contractile rates of contractile networks. Coarse-graining of these ensemble types from two sets of 16 motors on opposite ends of a stiff rod to two binders, each representing 16 motors, reduced force generation, contractility, and the total connectivity of filament networks for all ensemble types. A parallel cluster model (PCM) previously used to describe ensemble dynamics via statistical mechanics, allowed better contractility with coarse-graining, though connectivity was still markedly reduced for this ensemble type with coarse-graining. Together our results reveal substantial trade-offs associated with the process of coarse-graining NMMII ensembles and highlight the robustness of discretized catch-slip ensembles in modeling actomyosin networks.

**STATEMENT OF SIGNIFICANCE:** Agent-based simulations of contractile networks allow us to explore the mechanics of actomyosin contractility, which drives many cell shape changes including cytokinesis, the final step of cell division. Such simulations should be able to predict the mechanics and dynamics of non-muscle contractility, however recent work has highlighted a lack of consensus on how to best model the non-muscle myosin II. These ensembles of approximately 32 motors are the key components responsible for driving contractility. Here, we explored different methods for modeling non-muscle myosin II ensembles within the context of contractile actomyosin networks. We show that the level of coarse-graining and the choice of unbinding model used to model motor unbinding under load indeed has profound effects on contractile network dynamics.

## INTRODUCTION

Non-muscle myosin II (NMMII) is an actin-based motor that forms ensembles with 12-32 motor domain pairs extending from the ends of a rod formed by the bundled coiled-coil tails of the NMMII heavy chains (1–3). NMMII drives a myriad of cellular and tissue-level processes during cell migration, morphogenesis and cell division (4–7). Cytokinesis, one of these processes, is the last stage of cell division and is the physical division of one cell into two. Cytokinesis is powered by a cortical actomyosin band known as the cytokinetic ring, which forms in anaphase and constricts to invaginate the plasma membrane and partition the cytoplasm.

NMMII is capable of sliding actin filaments through its motoring activity (3, 8–11). However, NMMII motor ensembles are also robust F-actin crosslinkers (1, 8, 9, 12). Thus, it is unclear whether NMMII ensembles generate contractile forces through motor activity-dependent polarity sorting of actin filaments, which would induce filament buckling in a crosslinked network (13–16), or through passive crosslinking of depolymerizing or treadmilling F-actin (17, 18). The former hypothesis predicts complete failure of cytokinesis with motor-dead NMMII, while the latter predicts that motor activity is dispensable. Work with mammalian cultured cells and the budding yeast, *S. cerevisiae,* provided evidence for contractility with NMMII lacking the motor domain (17) or with motor-deficient NMMII (19), suggesting that NMMII primarily contributes as a crosslinker. However, work with similar NMMII mutants in *Drosophila* and *Caenorhabditis elegans* concluded that NMMII motor activity is required for cytokinesis in embryonic cells and tissues (20, 21). Separation of motor and crosslinking activities *in vivo* is confounded by observations that some motor-dead isoforms bind F-actin more tightly or for longer than wild-type NMMII and are thus crosslinking gain-of-function (20). Interestingly, contractile speed is not a linear function of the abundances of both NMMII and non-motor crosslinkers, as intermediate levels were observed to confer optimal contractile dynamics (15, 22–24). Defining the relative contributions of NMMII motoring and crosslinking to contractility is difficult, but nevertheless essential, since it will shed light on the mechanics of non-muscle contractility in the context of cellular processes such as contractility, cell motility and epithelial morphogenesis.

While biophysics, biochemistry and cell biology have revealed much about the contributions of NMMII to non-muscle contractility, the complex behavior of NMMII ensembles has remained hard to determine *in vivo.* A complementary approach can be found in agent-based modeling, wherein collective behaviors emerge from the explicit simulation of the components and their interactions. For this purpose, several open-source modeling suites that specialize in cytoskeletal dynamics are available (25–28). However, herein the question of how to best model ensembles of motors arises. Interestingly, the best method to model motor ensembles is unresolved. Even for single NMMII motors, unbinding dynamics under force is unknown. Generally, non-covalent bonds are described as slip bonds, where bond lifetime decays exponentially with increasing applied load (Fig. 1A). Some molecular motors, like dynein and kinesins, fall into this regime (29, 30); *in silico* unbinding dynamics akin to slip bonds can be modeled by use of Kramers’ theory or Bell’s Law (31). By contrast, some myosin motors or collections of myosin motors exhibit emergent catch-bond dynamics (32), behavior that can be modeled *in silico* by considering collections of motors (33, 34). Catch bond dynamics can be described generally by an exponential increase in bond lifetime as force is applied up to a critical point where the bond lifetime may plateau or eventually rupture under extreme force-load (Fig. 1B). Finally, atomic force microscopy performed with purified rat muscle myosin II heavy meromyosin (HMM) fragments (35) and optical tweezer experiments with single muscle myosin II ensembles (36, 37) have suggested that some myosin motor unbinding dynamics can be described by catch-slip bonds; which have been modeled within the context of motors by considering nucleotide-binding state (35). Catch-slip bonds exhibit a biphasic response to applied force; initially force-load increases the bond lifetime up to a critical point (the catch regime), after which the bond lifetime decreases (the slip regime) (Fig. 1C). While biological motors may exhibit complex force-dependent unbinding dynamics, these three simple scenarios cover the possible fundamental behaviors well. Nevertheless, the impact of unbinding dynamics on actomyosin network architecture, connectivity, and ultimately, contractility, remains unknown both *in vivo* and *in silico*.

**Figure 1.**
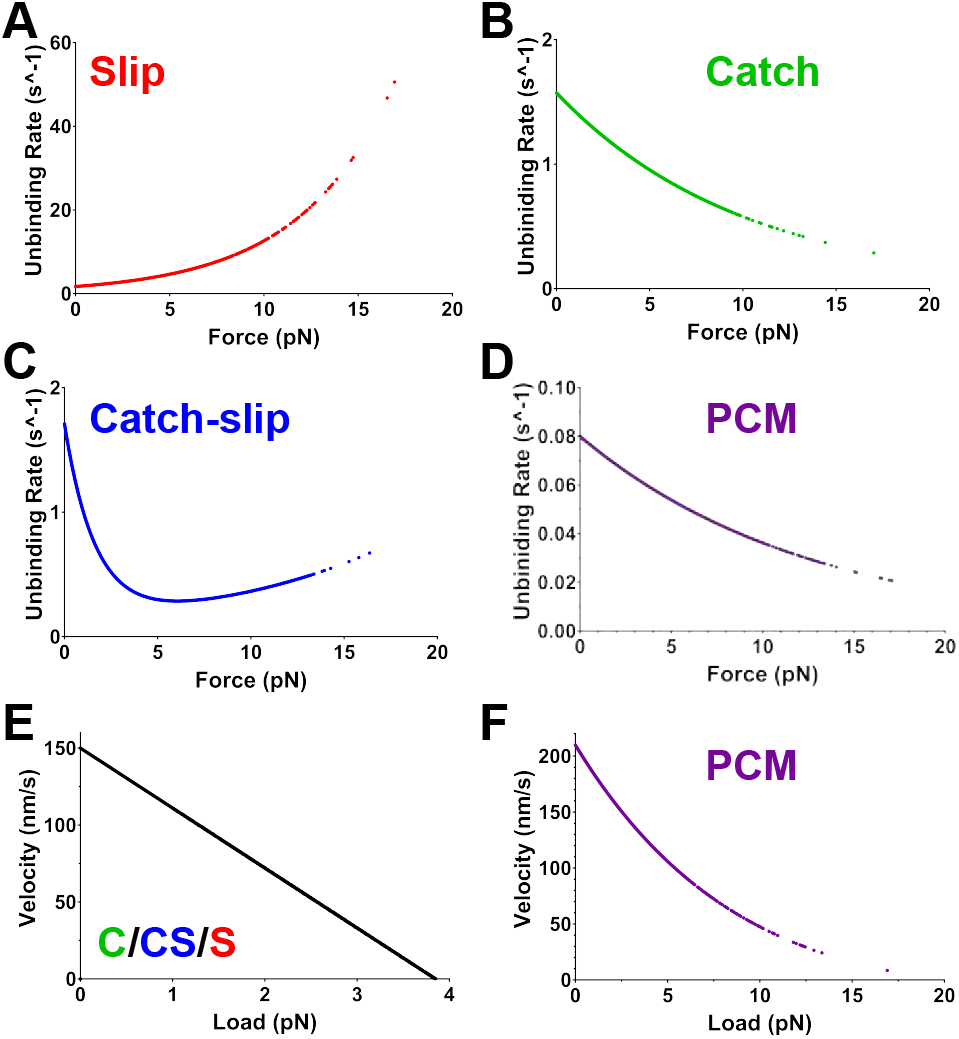
Single motor force-dependent dynamics. A-D) Different laws relating force to unbinding rates, used by Cytosim, and obtained here directly from the implementation. A) With slip bond dynamics unbinding rate exponentially increases with force. B) With catch bond dynamics, the unbinding rate decreases exponentially with force. C) Catch-slip bond dynamics show an initial catch-like behavior, here below ≈5 pN, and a slip-like behavior at higher force. D) The Parallel Cluster Model (PCM) exhibits catch bond dynamics with low unbinding rates. E) The force-velocity relationship for all but PCM bond types follows Eq. 10, where load and velocity are linearly related. F) The force-velocity relationship for PCM bond type, as characterized by Eq. 4, is exponential. Each curve is built from ~100,000 individual data points extracted from running simulations.

Published agent-based models of actomyosin networks have incorporated actin filament treadmilling (17, 18, 22), filament buckling (15, 16), varied crosslinker/motor ratios (15, 16, 22), and complex motor ensemble strategies (33, 34, 38–40). However, the relevance of these models to force generation in actomyosin networks is limited by how myosins were modeled. Specifically, many models employ simple dimeric processive motors some of which translocate in the opposite direction to NMMII (15, 16, 23, 40). In some cases motor ensembles were depicted either as multiple discrete binding entities (each representing one or a few motor heads) or as a single binding entity (coarse-grained; representing all the motors on one end of the filament). In the latter case, it is difficult to calculate aggregate binding and unbinding rates that correctly depict the behavior of multiple motors (22). The former is computationally intensive and may not realistically depict the complex crosstalk among motor subunits in an ensemble. Thus, there is no established solution for how to model the NMMII ensembles in an actomyosin system in such a way that optimizes simplicity and computational time, while recapitulating biological observations with maximal realism.

We sought to identify a suitable representation of NMMII that would recapitulate realistic motoring and crosslinking functions, as desired to simulate the mechanisms of non-muscle contractility. Using the agent-based modeling software, Cytosim (25) (www.cytosim.org), we compared several approaches to modeling NMMII motors in the context of contractile systems, consisting of actin-like filaments (referred to as “filaments” for simplicity), NMMII-like motor ensembles, and non-motor crosslinkers as necessary to general filament crosslinking and scaffold the network. Whereas previously we simulated NMMII filaments as simplified, coarse-grained, motor clusters represented by single actin binders on either side of stiff rods (22), here we compared such coarse-grained models with simulations of motor ensembles where motors were discretized and independent of each other. We also compared the behaviors of ensembles with motor domains exhibiting slip, catch, and catch-slip bond dynamics. We demonstrated that all three of these approximations can generate contractile networks, though they differ dramatically in their outcome, particularly in the macroscopic contraction rates and network order. We offer recommendations on motor modeling for optimization of calculation time, connectivity, and biological rigor.

## MATERIALS AND METHODS

### Cytosim base code

Cytosim, an Open Source Brownian motion C++ simulation program (25), was used to perform all agent-based simulations in this work. Within Cytosim, filaments are modeled as segmented lines, with bending elasticity as determined by the persistence length of F-actin, this is ~15 µm, resulting in a bending modulus of 0.06 pN·µm^2^. Filament movements are determined by an over-dampened Langevin equation:

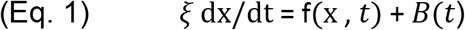

This equation accounts for system viscosity, drag (*ξ* as determined by the size of the filament) (41), thermal energy in term *B*(*t*), while f(x, *t*) includes the bending elasticity of the filament and the forces exerted by connectors (25). All simulations were initiated with an environmental viscosity of 1 Pa·s, similar to the viscosity of *C. elegans* zygote cytoplasm (42) and a thermal energy of 4.2 pN·nm. Actin dynamics were simulated as in previous models (17, 18), and include depolymerization primarily at pointed ends and polymerization primarily at barbed ends, with a net treadmilling rate of 0.1 µm/s and net depolymerization rate of 0.002 µm/s (43). Generic crosslinkers were modeled as rigid rods with actin and NMMII binding hands branching off them. Unless otherwise noted, all filament binders were modeled to have force-dependent unbinding rates. Unbinding dynamics of crosslinkers and scaffolds were modeled according to Kramers’ theory:

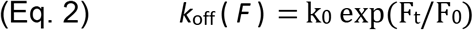

Here *k_0_* is the unloaded unbinding rate, *F_0_* is the characteristic unbinding force, and *F_t_* is the force felt by crosslinkers across their links at time *t*.

NMMII motor head unbinding dynamics were calculated by one of several methods which yield either slip bond, catch bond, or catch-slip bond dynamics. Motoring speed was calculated at each simulation frame by the formula: V_t_ =V_0_ (1-F_t_ / F_0_) (except for in the case of parallel cluster model (PCM) motors; where V_0_ is the unloaded motor speed, *F_0_* is the stall force, and *F_t_* is the load force, projected on the filament, at timepoint *t*. NMMII complexes were simulated as bipolar couples with actin-binding motoring domains attached at either end of a filament. Each actin-binding motor domain was modeled to simulate the dynamics of one or several NMMII motoring heads (between 1-16) for total NMMII valence of 12-32 motors (1–3).

The code of Cytosim was modified to enable new functionalities as follows:

1. Catch bond approximation using the Parallel Cluster Model (PCM):

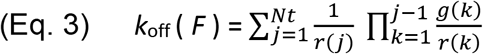

Where *Nt* is the number of motors, *r(j)* is the reverse rate of motor unbinding given j post-power stroke motors, *g(k)* is the forward rate of binding given *k* post-power stroke motors and *r(k)* is the reverse rate of unbinding given *k* post-power stroke motors; all of which are described in greater detail within their original work (33).
2. Bound-motor mean velocity approximation for PCM:

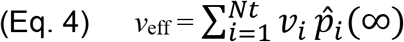

Where *v*_*i*_ is the complex velocity given *i* bound motors and *p̂*_*i*_(∞) the probability of an average *i* bound motors over infinite time (33).
3. Catch-slip bond approximation:

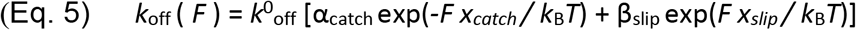

Where *k*^0^_off_ (or just *k_0_*) is the unbinding rate at rest, *F* is the force on motors, *x_catch_* and *x_slip_* are characteristic myosin II bond lengths, α_catch_ and β_slip_ are constants that weigh the catch and slip components, *k*_B_ is Boltzman’s constant, and *T* is temperature (34, 35).
4. Simple catch bond approximation:

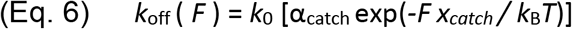

Which is a reduction of Eq. 5 where only the catch component is considered. General parameters used in most simulations are described and referenced in Table S1 of the Supporting Material.

### MM1: Single motor force-load simulations

(See Supplemental SINGLE_MOTOR configuration file for details). Simulations (Figure S1E of the Supporting Material; Movie 1) were set up to mimic laser trap force-load assays of single motors. A single filament with actin-like properties was immobilized in a horizontal orientation into the center of a periodic space 2 µm × 200 nm and subjected to a tug-of-war between a motor and a passive Hookean element. This passive element remained attached to the filament at all times. The motor was given a high binding rate to allow for more binding events (and therefore measurements) but its unbinding dynamic and motoring parameters followed exactly the different NMMII models. As these simulations run the motor pulls the filament leftward, loading the passive element and the motor alike. Simulations were run for 100,000 frames with 1 ms timesteps. At each step Cytosim output the load experienced by the motor along with its calculated unbinding rate and velocity. These data were plotted to visualize the effective force-load (Figure 1A-D) and force velocity (Figure 1E-F).

### MM2: Translocating ensemble simulations

(See Supplemental ENSEMBLE_TRANSLOCATION configuration file). To measure the movement of motor ensembles without interference from load and connectivity, we simulated motor ensembles, generated as monopolar complexes, that could bind to only one filament per binder at a time (Movie 2). Motors, either discrete or coarse-grained, were arrayed on one end of a stiff rod representing half of a NMMII ensemble. Immobile filaments 12.1 µm in length were placed horizontally, all parallel with right-pointing plus (barbed) ends. Motor ensembles were initially placed on the left side and allowed to translocate for 200 seconds along a periodic space 12 µm long. Ensemble movement speed was evaluated from positions recorded every 100ms and was calculated as follows: Positional information about all motor ensemble backbones (the “myosin fiber”) was pooled with information about the binding-state of all binders attached to each ensemble fiber backbone. Translocation speed was calculated for an ensemble only when the preceding time frame and the current time frame showed at least one of its ‘motors’ bound to an immobilized filament. Translocation was then calculated as the change in x-position (in nanometers) over time (each frame is 1 second apart). Average translocation was calculated per ensemble over the duration of the simulation. A population average and standard deviation were calculated for 200 ensembles of each type.

For visualization, kymographs were rendered in FIJI (ImageJ)(44). The simulation state was rendered every ten milliseconds, and the PNG images imported in FIJI. A 12 μm line was drawn over one of the immobilized filaments in each simulation and the ‘Multi Kymograph’ plugin was used to construct a kymograph for 300 timepoints from each simulation. Bound motor ensembles were visualized as grey dots, resulting in kymographs of streaks or lines moving from left to right (position) down the y-axis (time) of the generated images.

Motor ensemble binding percentage was calculated for all motors on all like-ensembles at each time point of the simulation, as well as for all ensembles as a unit (an ensemble was considered bound so long as one of its binders was actively bound to an immobilized filament). Averages and standard deviations were then calculated for each population of ensembles.

### MM3: Filament patch contractile system simulations

(See Supplemental FILAMENT_PATCH configuration file). The simulations (Figure 3B; S2; Movie 3) generated filament patches in a two-dimensional circular space the size of a fission yeast cell (radius of 1.5 μm) and seeded with motors and crosslinkers. Based on estimates for fission yeast contractile rings (45, 46), we seeded 360 dynamic actin-like filaments of length 1.0 ± 0.3 μm, which depolymerize at a rate of 2 nm/s, in random orientations within the circular space. 360 motor ensembles and variable numbers of crosslinkers, from 1,000 to 12,000, were then added and allowed to bind filaments before the simulations were run. We used a circle-fitting method (described below) to estimate the contractile rate in terms of a radial speed in nm/s. Compression force (contraction) was estimated in these patches along virtual planar sections, rotated over 360 degrees in increments of 10 degrees. At each plane, Cytosim reports the tension/compression across all fibers intercepting the plane. The force reported is the average of the sum total force experienced by actin-like filaments in all simulations for each set of simulations. Maximal radial contractile speed was reported for all individual simulations of the same ensemble type but with varying amounts of crosslinkers. The number of crosslinkers allowing for maximal patch contractility was that which led to the highest maximum contractile speed for all simulations of like ensembles. Five simulations were then run with the amount of crosslinker generating peak contractility for each ensemble type and total network connectivity was calculated for these contractile networks (See MM7).

### MM4: Periodic linear contractile system simulations

(See Supplemental LINEAR_NETWORK configuration file). The simulations shown in Fig. 4 used periodic boundary conditions to prohibit the network from shortening, to estimate the force produced by the contractile network. The 2D simulation space was periodic in the X-direction but not in Y with dimensions that roughly match the fission yeast contractile ring at the onset of contractility: length of 9.42 µm, 360 actin filaments, 360 motor ensembles, and 1,000-3,000 crosslinkers (45, 46). The amount of crosslinker was selected based on peak contractility curves generated in patch network analysis (see MM3). Periodic simulations were run for 400 seconds of simulated time, during which time most linear networks ruptured, marked by the formation of a filament-free gap and recoil of the network into clusters. Periodic network simulations were analyzed for several features including occurrence of failure (rupture), tension before network failure, compression after network failure, and number of motors bound. These analyses were performed by processing Cytosim reports in MATLAB. Briefly, a plane parallel to the Y-axis was scanned, with 100 nm steps, across the simulated space at each simulation time point. As before, force felt across actin-like filaments (either compression or tension) was summed up at each time point to provide a network-wide output. Maximal tensile force was reported before network failure (when networks are not contractile yet); maximal compressive force was reported after failure when networks collapse into clusters of filaments. Rupture was detected when no intersecting filaments were reported for at least one plane.

### MM5: Contractile ring simulations

(See Supplemental CONTRACTILE_RING configuration file). The simulations shown on Fig. 6 (Movie 5) of a 2D contractile ring were set up in a circular space with a radius of 1.5 µm. Environmental parameters were as in our actin walking simulations. At initiation, the space was populated with 360 actin filaments of 1 ± 0.3 μm long, and 5000 generalized crosslinkers. Actin filaments were placed within 60 nm of a circle of radius 1.490 µm, tangent to the circle. The crosslinkers were placed just overlapping and outside of the actin filaments. The simulation was allowed to generate binding events without moving the filaments for two seconds, and then 360 myosin ensembles were added to the ring space. Following another two seconds during which only binding and unbinding could occur, the Brownian dynamics were enabled, and run for 400 seconds. For simplicity, the abundance of components was constant throughout the simulations. However, unbound components could diffuse far enough away from the ring to never reincorporate, and thus be non-productive. Actin filaments were dynamically treadmilling with net polymerization at one end and net depolymerization at the other (0.002 µm/s). For more details, see the sample code.

### MM6: Circle-fitting of contractile patch simulations

Patch and ring network (MM4 and MM5) closure dynamics were quantified by estimating the speed of radial change. Given the variable nature of these networks in our simulations, an average radius R was first calculated for all time points. From Cytosim, we first exported coordinates for the centers of all myosin ensembles at each time point as .CSV files. These CSV files were reorganized for readability then ported as tabular data into MATLAB (MathWorks) as structures consisting of coordinate matrices for each frame of simulated data.

The Pratt method (47) was then used to fit a circle to each set of coordinate points representing the network, allowing us to estimate *R(t)*, from which we derived raw radial speed, *S*:

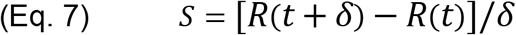

Where radius is recorded in µm, and *δ* is the timestep in second. We also used the radius measurements to calculate network closure percentages, *C*, as:

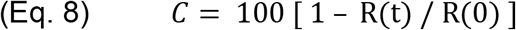

The contractile patch network perimeter was also calculated for each frame. Simulated networks frequently constricted in in non-symmetrical and off-center shapes. As such, we reduced fitting-error noise by performing a moving average of speed, radius and perimeter data with a window of five frames. We also controlled for the possibility of a network closing partway and then reopening after losing connectivity or compressive forces. This was done by calculating the derivative of the closure percentage and finding values that were smaller than −1 %·min^−2^ suggesting significant relaxation of the network. If significant relaxation was recorded at frame *n* data from frames *1 to n-1* were kept for calculation of speed while all other data were discarded. Similarly, to prevent noise from data that occurred after full network constriction, we discarded data from beyond 95% closure.

Once all data were smoothened for each individual simulation, we combined the datasets for all like simulations and calculated average and standard deviation of speed, percent closure and perimeter. Average speed data were smoothened via a five-frame window moving average, as before (22).

### MM7: Network Connectivity Estimations

We extended Cytosim report functions to export meaningful connectivity data specifically for this project. First, to report motor ensemble-based connectivity (λ), we coded in a function that found all motor subunits from a myosin ensemble and reported the unique identifier of all associated filaments (one filament per attached motor). These data, in CSV format, were then ported to MATLAB where a simple function detected the number of unique filaments interacting with each motor ensemble at each time point. The interaction number was then averaged for all motor ensembles throughout the simulation or up until 80% closure was achieved. The average number of unique filaments attached to an ensemble, Γ, was then used to calculate its contribution to connectivity:

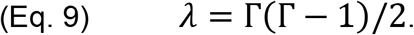

The λ for all motor ensembles were summed to calculate the final connectivity provided by NMMII ensembles. The same approach described above was used to sum connectivity of non-motor crosslinkers. The different connectivity numbers were summed to calculate the total network connectivity in these simulations.

### MM8: Statistical analysis of data

All replicate data were reported as a mean ± one standard deviation from the mean. Comparison of mean data between different bond type and ensemble discretization or coarse-graining was performed by use of unpaired t test analysis in GraphPad Prism8 (www.graphpad.com).

### MM9: Coarse-graining of non-PCM motor ensembles

Non-PCM motor ensembles were coarse-grained fully such that all 16 motor components on either side of bipolar filaments were collectively represented by a single filament binder. To maximize the behavioral similarity between coarse-grained and discretized motor entities, we performed parameter sweeps testing a range of values of the binding rate, unbinding rate, unloaded speed, and stall force of each motor type in immobilized filament translocation simulations (see MM2). As described above, ensemble translocation speed and ensemble binding ability served as readouts for the performance of coarse-grained ensembles to the original discretized ensembles of the same bond type. 250 simulations were generated by sweeping the four aforementioned parameters and comparing average behavior of 40 ensembles per each simulation to the established behavior of the discrete ensembles of the same bond type. Simulations that minimized the difference in average ensemble behavior were rerun 4 additional times to generate average data for 200 total ensembles, which was then statistically compared to the data from 200 discrete ensembles of the same bond type to test for indistinguishable performance. Parameters were selected only if the comparison of the average ensemble translocation speed and average binding ability both resulted in insignificant difference (p values above 0.2 were selected as a threshold).

## RESULTS

### Different bond types generate different force-binding curves in silico

By default, Cytosim filament-binders follow Bell’s Law and Kramers’ theory (31) to calculate force-dependent unbinding of components; unbinding rate increases exponentially as force is applied to filament binders, as seen for biological slip bonds. Since myosins have been observed to exhibit catch-and catch-slip bond behavior (32, 35), we first modified Cytosim base code to include models for the force-dependent regimes of catch bonds and catch-slip bonds that can be designated for any class of filament-binders. At each simulated timestep, forces were calculated and used to estimate the unbinding rate of the individual components based on which method was selected. To verify our implementation, we first set up simulations to mimic *in vitro* single motor laser trap experiments (see Methods MM1). Briefly, single simulated motors pulled against a passive Hookean spring linker, load on the motors was varied, and unbinding rate was reported. Importantly, motors unbinding according to the three different models performed markedly different from one another (Figure 1A-C). We compared these unbinding dynamics to those of the previously published Parallel Cluster Model (PCM), which resulted in emergent catch-bond dynamics (Figure 1D), but with an unbinding rate roughly 20-fold lower than that observed with any of the other models. Thus, bond types other than simple slip-bonds can now be implemented in Cytosim.

### PCM motors have different force-velocity profiles

The effect of force on velocity in Cytosim can be calculated by converting force exerted on the motor into load; which is assistive if directed in the direction of motor translocation, and resistive otherwise. Displacement (*D*) is then calculated as:

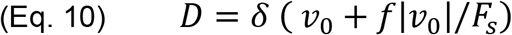

Where *v*_0_ is the unloaded speed, *f* is the directional load, and *F*_*s*_ is the stall force of the motor. This results in linear force-velocity curves for single motors (Figure 1E, Figure S1 in the Supporting Material) of the catch, slip and catch-slip types.

However, the PCM approach includes an independent way of calculating an average effective velocity, *v*_*eff*_, that considers the average behavior of a binder with *i* bound motors (Eq. 4). Since *v*_*eff*_ is calculated as the average behavior of a stochastic ensemble, force is considered as a positive net value instead of being treated as directional load on the motor ensemble. The result is a force-velocity curve with an exponential drop-off, which reaches zero velocity closer to 20 pN of force per motor (Figure 1F), compared with a linear relationship with the same derivative at zero (Figure 1E). Both methods of calculating single motor load-dependent velocity result in curves with characteristic behaviors comparable to measurements made *in vitro* for muscle myosin and *in vivo* for motors (48, 49). Thus, *in silico* motors can recapitulate force-dependent behavior demonstrated *in vitro*.

### Discrete ensembles translocate on immobilized filaments

We next built motor ensembles akin to structures reported from *in vitro* reconstituted mammalian NMMII (1–3) and tested them in translocation assays *in silico*. Ensembles of each motor type were constructed by permanently coupling motor agents to one end of a long stiff rod. Ensembles were generated in a monopolar formation, unlike bipolar ensembles *in vitro*, to simplify translocation simulations and reduce the potential of ensembles interacting with and binding to multiple filaments at once. Fully discretized monopolar versions of each ensemble were constructed by attaching 16 motor agents to one end of a stiff rod (Figure 2A).

**Figure 2.**
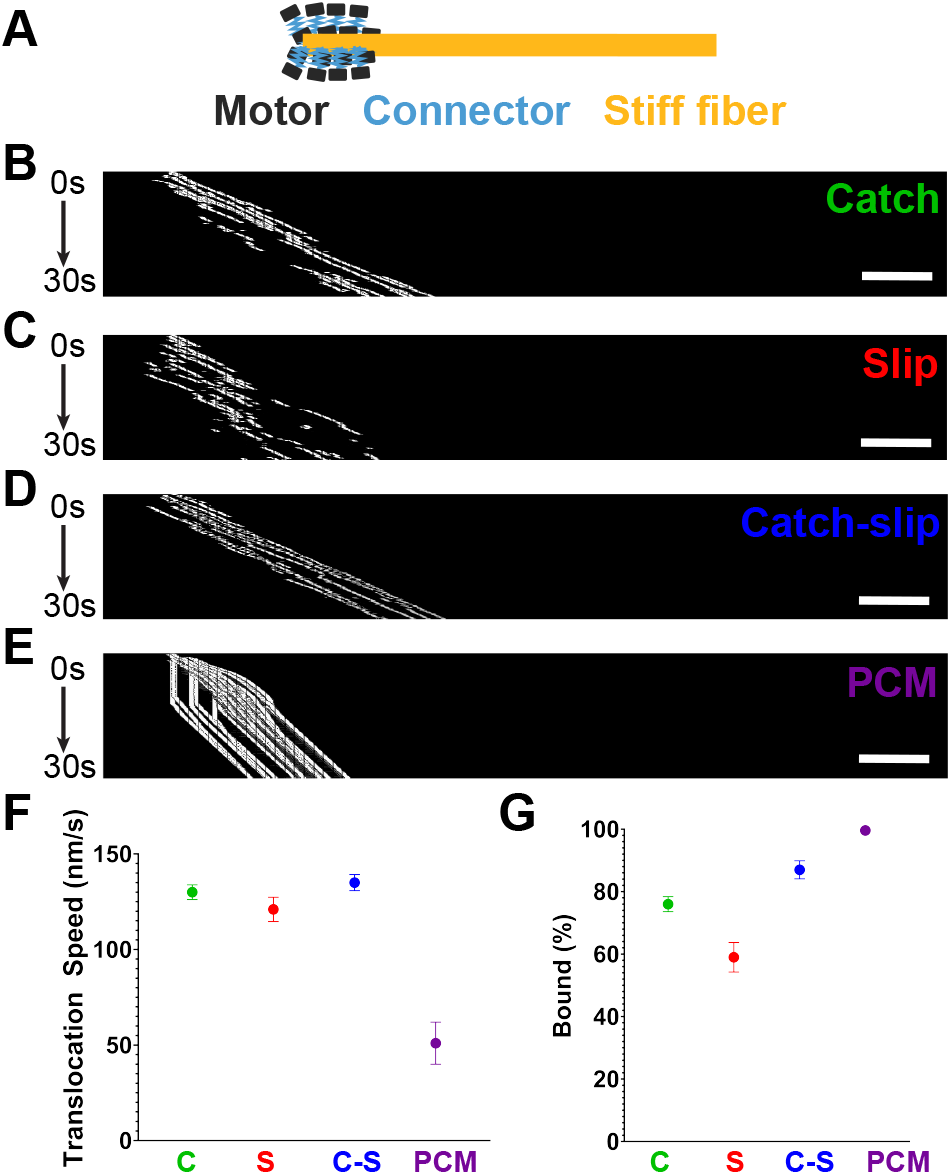
Discretized motor ensemble translocation on immobilized filaments. A) Schematic representation of a monopolar motor ensemble where individual NMMII-like motors (grey) are affixed to one end of a stiff rod representing NMMII bundled tails (yellow), by springs (blue). B-E) Kymographs of catch bond (B), slip bond (C), catch-slip bond (D) and PCM (E) ensembles (grey) translocating on immobilized filaments (not shown). All kymographs extend over 12 µm (x-axis) and 30s (300 frames) in duration (y-axis). F) Translocation speeds for each ensemble type shown in B-E. G) Binding percentage (over total time) for each ensemble type shown in B-E. Points show mean; bars show standard deviation. Sample size is 5 simulations with 40 ensembles each for a total of 200 data points for each case. Scale bars are 1 µm.

We first simulated discretized ensemble translocation on immobilized filaments for all three bond types and the PCM model (Movie 2). Catch and catch-slip bonding ensembles motored with similar, though statistically distinguishable speeds of 130±4 nm/s (Figure 2B, F) and 135±4 nm/s (Figure 2D, F) respectively. Both of these translocation speeds fall in the reported range of ~133±75 nm/s for *in vitro* reconstituted NMMIIA ensembles (1). Slip bonding motor ensembles also translocate well, though at 121±6 nm/s (Figure 2C, F), significantly slower than expected. Finally, PCM discretized motors translocate even more slowly at 51±11 nm/s (Figure 2E, F). *In vitro* reconstituted NMMIIA ensembles were previously shown to be highly processive, remaining bound to actin fibers through most of their translocation time (1). Thus, we set out to calculate the average percent of time bound for each type of motor ensemble and found that PCM ensembles were the best binders, remaining bound 99.6±0.41% of total simulation time. Catch-slip bond ensembles were second best at binding, remaining bound 87±2.88% of total simulation time. Catch and slip bond ensembles were markedly worse binders with 76±4.85% and 59±4.74% binding percentages respectively (Figure 2G; Table S2 of the Supporting Material). Thus, while all motor bond types were capable of binding to and translocating on filaments, only catch and catch-slip bond ensembles appeared to recapitulate established behavior of reconstituted NMMII ensembles.

### Catch-slip bond ensembles contract patch networks fastest

After establishing that motors with all bond types can translocate under low load, we next characterized how different motor ensembles behave in contractile networks. For these more physiologically relevant networks, we simulated bipolar ensembles with 16 discretized motors on either end of the stiff rod (1–3; Figure 3A). Bipolar motor ensembles were then seeded into simulations bearing randomly oriented actin-like filaments, which treadmill and depolymerize over time, and generic crosslinkers that can bind motor ensemble rods and actin-like filaments, similar to anillin proteins (50). Based on measurements from fission yeast cytokinetic rings (35), motor ensembles and filaments were present at a 1:1 ratio (360 of each), and 1,000-12,000 crosslinkers were included. Varying crosslinker amount affected the maximum contractile speed for all three bond types, including PCM catch bonds (Figure 3B; Figure S2 of the Supporting Material). Interestingly, only catch-slip and PCM motor ensembles exhibited a peak in contractility at intermediate crosslinker concentration (within the presented range) as predicted from cell biological, *in vitro* reconstitution, and earlier *in silico* work (15, 16, 22, 23; Figure 3C), which was around 3,000 crosslinkers or ~8-fold more crosslinker than motor ensembles per fiber for catch-slip, and around 2,000 crosslinkers or ~5-fold more crosslinker than motors per fiber for PCM. Catch and slip bond ensembles revealed optimal contractility with lower connectivity when crosslinker sweeps were repeated with 200-1,200, both peaking around 1,000 crosslinkers (Figure S3A of the Supporting Material), or ~3-fold more crosslinkers than motors per fiber. When we compared what crosslinker level allowed maximum contraction speed for all bond types, we saw that all but the PCM model exhibited the same initial behavior up to ~1,000 crosslinkers. After this point, slip and catch bond ensembles appeared to lose productive force generation and dropped off in contractile speed with peaks at 87±3.97 nm/s for catch bond and 70±4.97 nm/s for slip bond ensembles. Catch-slip ensembles, in contrast, peaked at 106±2.97 nm/s. PCM ensembles were the least effective force generators with peak contractile rates of 8±0.37 nm/s (Figure 3C).

**Figure 3.**
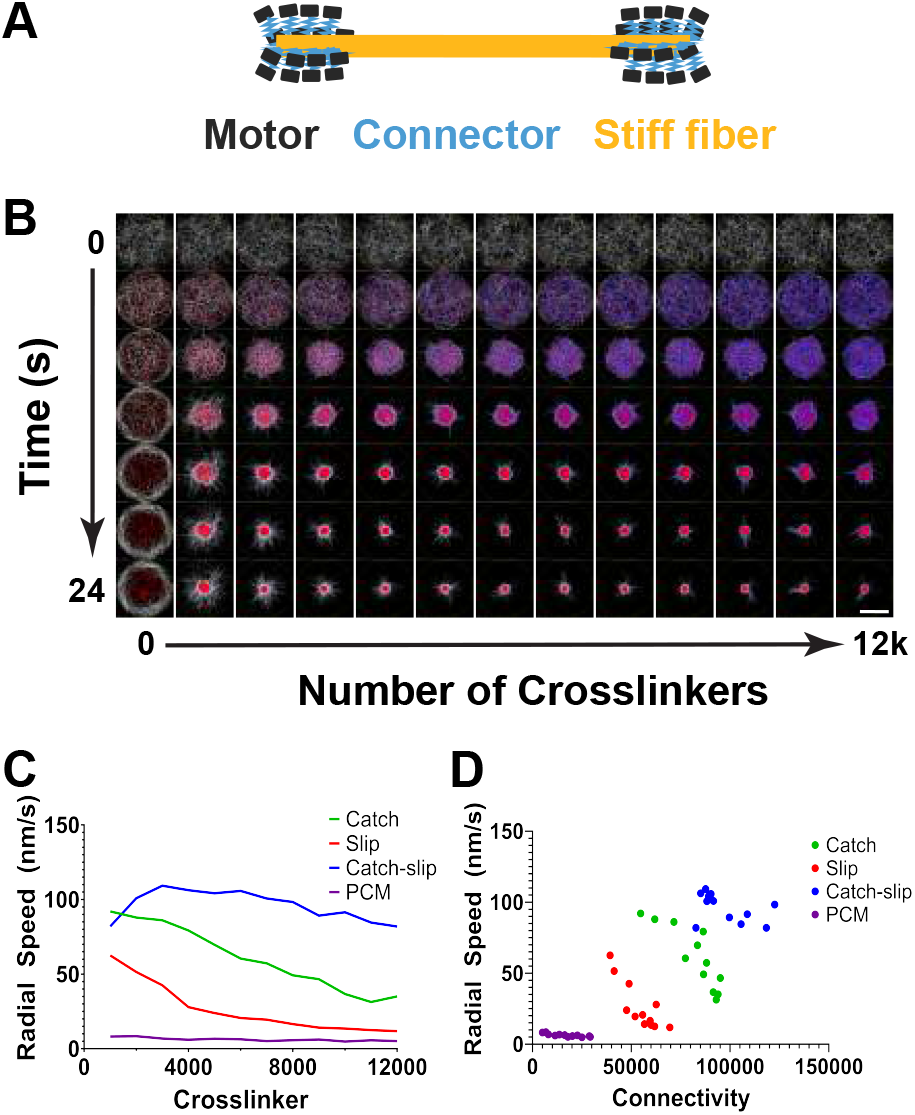
Contractile patch simulations of discretized motor ensembles. Schematic representation of a bipolar motor ensemble where individual NMMII-like motors (grey) are affixed to both ends of a stiff rod representing NMMII bundled tails (yellow), by springs (blue). B) Representative simulations of contractile patches. Actin-like filaments (white) are seeded with motor ensembles (red) and varying amount of crosslinkers (blue) ranging from 0 to 12,000 (along the x-axis). Simulation snapshots are shown from top (0s) to bottom (24 s) with 4 s intervals. C) Radial closure speed (in nm/s) of contractile patches for each ensemble type as a function of crosslinker amount. D) Radial speeds for all conditions of each ensemble type (dots) as a function of calculated total connectivity (see Methods MM7) of the patch networks. Scale bar (bottom right) is 1.5 µm.

We next compared the effect of total connectivity, calculated as the sum connectivity contributions of motor ensembles and crosslinkers (see Methods MM7). Similarly, catch-slip bonding ensembles were able to generate productive force with higher connectivity than ensembles with either of the other bond types (Figure 3D). Maximal contractility occurs for catch-slip bond ensembles with connectivity of 79,000±4,900, at 59,000±7,200 for catch bond ensembles, at 41,000±9,200 for slip bond ensembles, and at 6,900±120 for PCM ensembles. Thus, PCM ensembles do not confer contractility even in highly connected networks, despite being better at binding in translocation simulations. Interestingly, discretized PCM ensembles do not confer much connectivity themselves, but were robust bundlers unlike other ensemble types, as evidenced by significant filament alignment not seen for other ensemble simulations (Figure S3B) and quantified by network directionality analysis which revealed peak order of ~0.6 for PCM ensembles and ~0.37-0.47 for all others (Figure S3C-E; Movie 6). Thus, even in these simplified contractile systems we saw different contractile dynamics with all four different motor bond models.

### Motor ensemble dynamics on linear actin networks

The stark contrast among motor bond types on contracting patches suggested that these variably-binding ensembles would also achieve different levels of tension. To characterize the effect of motor bond type on tension generation, we constructed periodic linear networks that generate tension and cannot contract. We simulated linear bundles the length of the *S. pombe* contractile ring circumference, in a 2D space with periodic horizontal boundaries. As with the patch simulations, we modeled 360 actin-like filaments that treadmill and depolymerize over time, with 360 motor ensembles. Crosslinker abundance was based on that which allowed maximum contraction speed in patch simulations. In these linear network simulations, mounting tension was marked by a thinning of the filament band. Ultimately, all linear networks simulated with slip bonds ruptured (n = 20), as did simulations with catch and catch-slip bond ensembles, but rupture only occurred in 12 out of the 20 PCM simulations (Figure 4A; Movie 4), marked by a gradual emergence of compressive force (Figure S4A). As with actin patch simulations, catch-slip bond ensembles generated the greatest global compression force, which was several-fold higher than that of the other discretized ensemble simulations (Figure 4B). In addition, a higher percent of catch-slip bond ensembles was bound, than for all other ensemble types (Figure 4C). Interestingly, only ~30% of PCM ensembles were bound. This can be explained in part by the overall lower force generated by linear networks bearing PCM ensembles, which was ~1,000 pN before network failure and -60 pN after, 10-30-fold lower than that reported for catch-slip bond ensembles at these same points. PCM ensembles, which behave as catch bonds, are poor binders unless significant force is applied to them, here these ensembles felt too little force to remain effectively bound.

**Figure 4.**
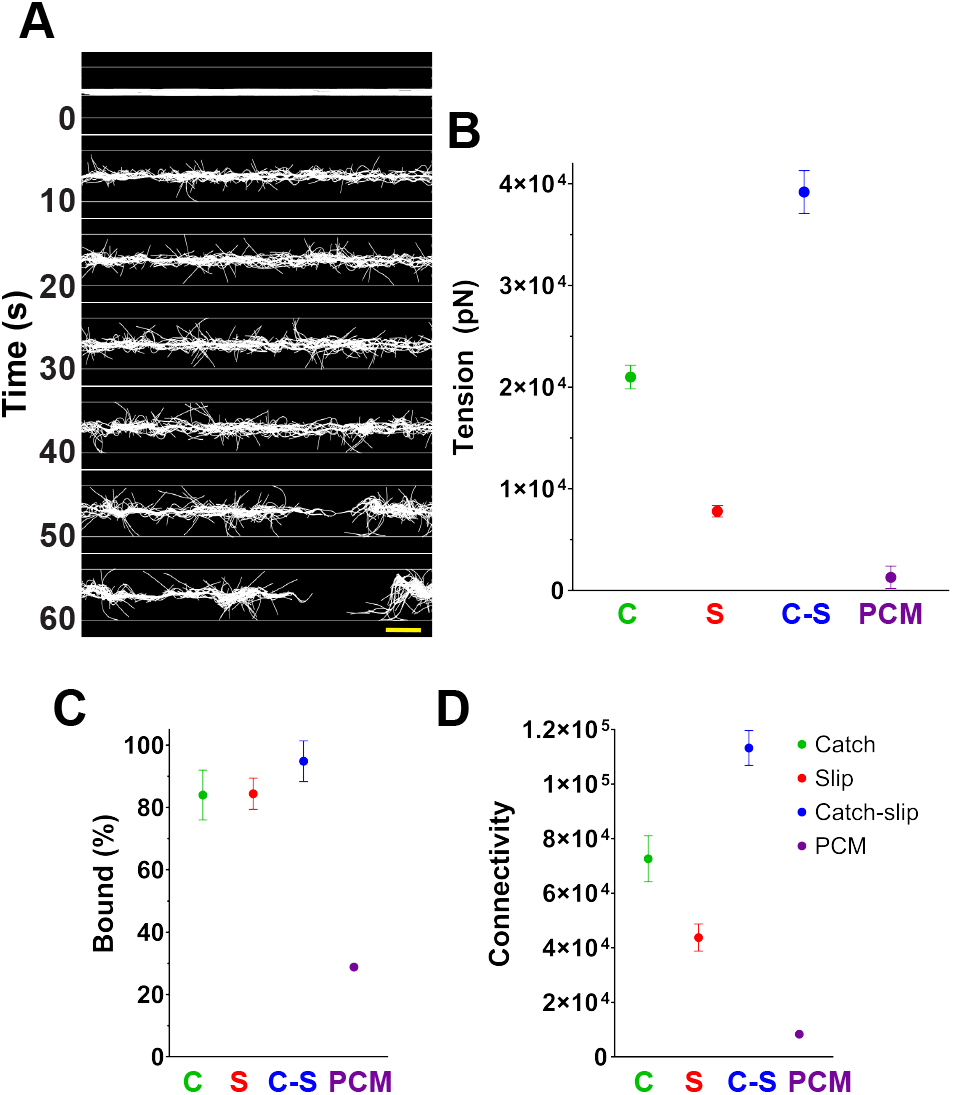
Linear periodic contractile network simulations of discretized motor ensembles. A) Representative simulation of a linear network that is closed on itself across periodic boundaries. Actin-like filaments (white) are initially placed horizontally but in random left/right orientation and position. Motors (not shown) exerting tension eventually lead to rupture (at t=50 s). Scale bar (yellow) is 1 μm. B) Average tension (before rupture) through a cross-section of the linear networks with each of the ensemble types. C) Percentages of bound motors (before rupture) for each ensemble type. D) Average total connectivity (see Method MM7) before rupture, for each of the motor types. Points show mean; bars show standard deviation. Sample size is 20 simulations for each motor ensemble type.

We also assessed network connectivity conferred by crosslinkers and motor ensembles. Similar to their behavior on contractile patches, catch-slip bond ensembles in bundles generated the highest total connectivity, topping out at ~110,000, and PCM ensembles were the lowest, topping out at ~8,000 (Figure 4D). Connectivity across all simulations is higher than the connectivity for the same discrete ensemble type in actin patches, suggesting that network architecture also contributed to total connectivity.

### Coarse-grained PCM motor ensembles generate more force than discretized PCM ensembles

Modeling motor ensembles with discretized motor domains takes full advantage of the powers of agent-based modeling but is computationally costly. Indeed, NMMII ensembles have conventionally been fully or partially coarse-grained in agent-based models (22, 33, 39). We therefore next set out to compare fully coarse-grained ensembles with the different bond types to their discretized counterparts in our simulation setups. For PCM ensembles, this was done by changing a single parameter representing ensemble size that dictates the number of motors being estimated as a single entity by statistical mechanics (33). Increasing the coarse-graining of PCM ensembles (while keeping the total number of motors depicted at 16 per side) significantly increased translocation speed and processivity (Figure 5A). Filament patches with coarse-grained PCM ensembles had ~3-fold lower connectivity and 13-fold higher contractile speed than patches with discretized PCM ensembles (Table S3 of the Supporting Material). Similarly, linear actin networks with coarse-grained PCM ensembles generated 10 times as much force as simulations with discretized PCM ensembles and always (n = 20) ruptured within 9±2.86 seconds of simulated time compared to only 12 out of 20 that ruptured at 382±20.15 seconds for discretized motors. In addition, a greater proportion of coarse-grained PCM ensembles were bound, compared to discretized ensembles (98.1±0.43% and 28.8±0.16%, respectively; Table S4 of the Supporting Material), which is not surprising given the higher force produced by networks with coarse-grained ensembles.

**Figure 5.**
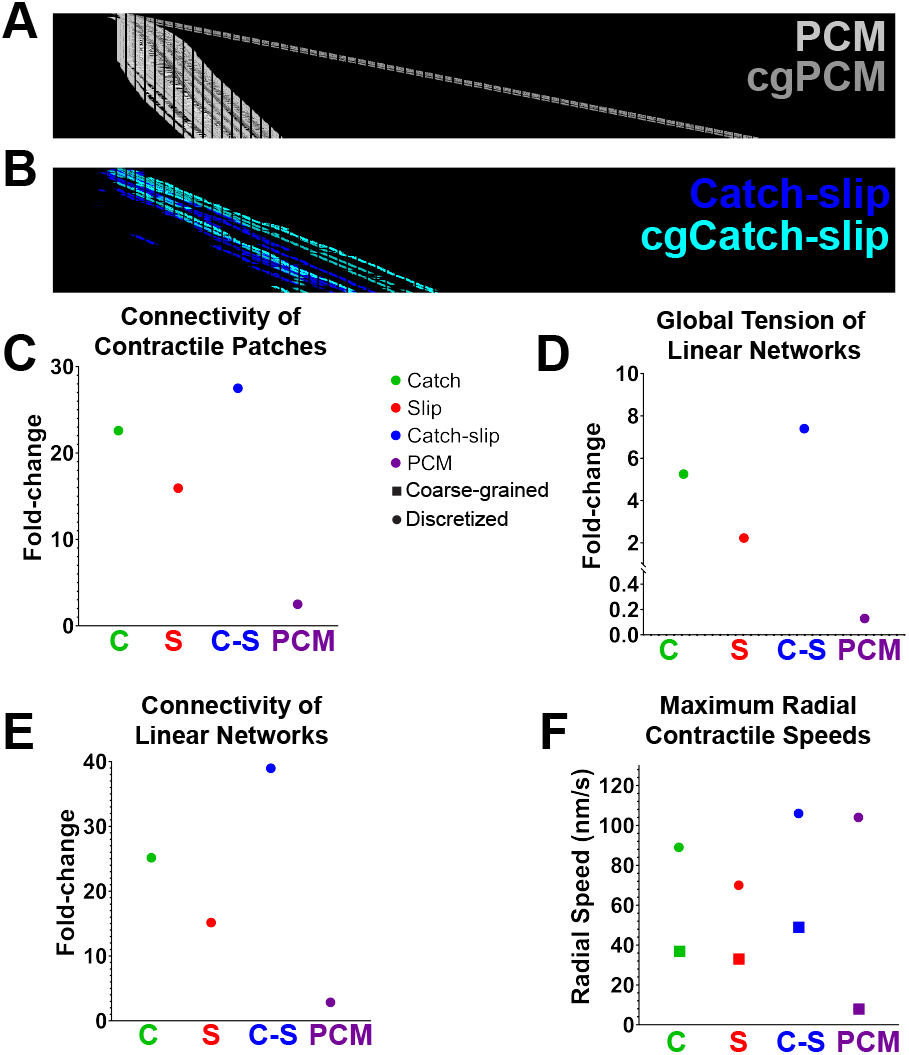
Comparison of discretized and coarse-grained motor ensembles. A) Kymograph comparing discrete (light grey) and coarse-grained (dark grey) PCM ensembles translocating on immobilized filaments. B) Kymograph of discrete (dark blue) versus coarse-grained (light blue) catch-slip ensembles showing similar kinematics. Kymographs are 12 µm (x-axis) by 30 s (300 frames; y-axis). C) Fold-change in global tension for contractile patches with discrete versus coarse-grained ensembles of all types. D) Fold-change in global tension for periodic linear networks with discrete versus coarse-grained ensembles of all types. E) Fold-change in Binding % on linear networks between discrete and coarse-grained ensembles of all types. Fold-changes under 1 indicate fold decrease from discretized to coarse-grained ensembles. F) Maximal radial contractile speed versus connectivity for all ensemble types. Squares show speeds for contractile networks with discretized ensembles; circles show speeds for networks with coarse-grained ensembles. Sample size is 5 simulations for all data. Standard deviations in F are too small to visualize. Scale bars are 1 µm.

For the PCM method, some changes in contractile dynamics arise due to different binding and motoring ability conferred by the averaging of collective motor behavior. However, as expected from the representation of fewer filament-binding sites on coarse-grained motor ensembles, in both patch simulations and linear actin network simulations, coarse-grained PCM ensembles conferred lower total network connectivity compared to discretized PCM ensembles. Therefore, connectivity could also explain some of the differences in contractile dynamics.

### Coarse-grained ensembles of non-PCM motors achieve less contractility

We expected that coarse-graining of all motor ensemble types would generally reduce network connectivity due to a decrease in the number of available filament-binders per ensemble. By generating coarse-grained ensembles with binding and motoring behavior comparable to discrete ensembles with the same motor type, we could test whether the changes in connectivity resulting from coarse-graining contributed significantly to changes in ensemble and network dynamics. Thus, we next set out to generate coarse-grained ensembles of catch, slip, and catch-slip bond types. These methods require parameter values which have not yet been measured for aggregated motors. Thus, we first used immobilized filament ensemble translocation simulations (see Methods MM2) to test a range of values for several parameters to establish the coarse-grained versions of ensembles for each bond type that translocated similarly to discretized ensembles of the same bond type. Binding and unbinding rates, as well as unloaded speed and stall force, were all swept in translocation simulations in order to establish the parameters for ensembles with translocation speed and percent bound outputs indistinguishable from the discretized ensembles of the same motor type (Figure S4B) (see methods in MM9). Qualitatively, coarse-grained ensembles exhibited similar dynamics to their discretized counterparts (Figure 5A, B; Movie 7). Statistical analysis revealed no difference within coarse-grained and discretized pairs, in terms of either translocation speed or ensemble binding percentages (Table S2l).

Coarse-grained ensembles seeded onto filament patches conferred lower connectivity than the discretized counterparts for each bond type: 15-fold decrease in connectivity for slip bond ensembles, 20-fold decrease for catch bond ensembles, and 30-fold decrease for catch-slip bonding ensembles at peak contractile speeds (Figure 5C). This decrease in connectivity occurs in parallel with a ~2-2.5-fold drop in contractile speeds for all three bond types (Figure 5F). Linear networks with coarse-grained ensembles of the three types showed similar drops in connectivity (Figure 5E), and force generation (Figure 5D), which resulted in delayed rupture. Given the artificially high binding rates and low force-dependent unbinding rates of coarse-grained ensembles, it is not surprising that all three bond types exhibited high ensemble bound percentages. Coarse-grained ensembles with catch and catch-slip bonds were virtually indistinguishable at 98.9±1.5% and 98.2±1.2%, with slip ensembles not much lower at 94.6±1.1% (Table S4). In sum, coarse-graining of all non-PCM ensembles resulted in a reduction in connectivity, as expected, along with drops in force generation and contractile speed.

## DISCUSSION

### Summary

We compared the behavior of NMMII-like motor ensembles using four modeling strategies covering three unbinding behaviors; catch, catch-slip, and slip bonds. We further compared the effects of modeling ensembles of all four strategies as either fully discretized entities with individual motor agents or as fully coarse-grained entities leaving only two motoring agents per NMMII ensemble, behaving as a collection of motors. We showed that all unbinding laws affect motor binding and translocation speeds as well as network properties such as contractile speed and connectivity. Importantly, coarse-graining of ensembles significantly alters network connectivity and global contractile rates independently of an effect on individual motor speed and binding dynamics for all four models.

### PCM ensembles

As previously reported (33), catch-bond behavior is an emergent property of motors operating according to the parallel cluster model. Here we expand this observation to the emergence of catch dynamics for ensembles of 32 motors, whether motors are fully coarse-grained or fully discretized. However, coarse-graining does alter the binding time both at the individual agent level and at the ensemble level. PCM ensembles can generate more force as coarse-graining increases due to an increase in the number of bound motors and therefore an increase in the average effective velocity. Due to a higher force generation, the catch-bond behavior of coarse-grained PCM ensembles ensures that they are better binders in both patch and periodic filament network simulations. Compared to all other motor strategies, discretized PCM ensembles are the least effective force-generators but coarse-grained PCM ensembles are among the best at force generation (only catch-slip ensembles are better).

Two caveats exist here, however. First, PCM motors were simulated using parameters estimated from muscle myosins which are known to have different biophysical properties than non-muscle myosins. Future work should therefore carefully adapt the PCM method with NMMII parameters. This is important, especially given that all other motor types herein were simulated using parameters estimated for NMMIIA previously (34). Second, network connectivity, which has been repeatedly shown to be important for contractile dynamics (15, 16, 22, 23), is lower for both modes of PCM ensemble simulation. In the case of the more processive coarse-grained PCM ensembles, the network connectivity is decreased because each ensemble can bind a maximum of two actin-like filaments instead of ~20-30. While this may be compensated *in silico* by adding more non-motor crosslinkers, we lack the ability to discern the direct effect of motor-based connectivity on contractile dynamics. We therefore suggest that the coarse-grained PCM method is ideal for fast and robust simulation of large processive motor ensembles so long as questions about connectivity need not be addressed.

### Slip bonding motor ensembles

Slip bond motor ensembles are much poorer force generators than PCM, catch or catch-slip bonding ensembles with the same level of coarse-graining. This result is not surprising given the inherent decrease in binding lifetime as a contractile network comes under compression force. Fully discrete slip ensembles are fast motors when they are bound, largely due to their quick unbinding that likely often happens before the motor reaches stall force, but the same characteristic makes these motors such poor binders that they exhibit little processivity in our immobilized filament simulations. At very low ratios of non-motor crosslinkers to motor ensembles, slip bond ensembles can produce contractile speeds around 70 nm/s with peak force around 600 pN across actin patches. Coarse-graining slip bond ensembles results in a significant drop in all metrics except force generation (excluding unloaded binding and translocation speed, which were matched to their discretized equivalent). In the case of slip bond ensembles, coarse-graining does not appear to provide any benefit beyond simplifying the structure of motor ensembles and reducing computation time. Use of slip bond motors in simulations would therefore be best suited to modeling single motors that follows such a law.

### Catch bonding motor ensembles

Catch bond motor ensembles are predicted to be good crosslinkers given their inherent increase in binding lifetime as force is applied. In practice, however, we see comparable binding percentage for discrete catch bond ensembles as for slip bond ensembles (84±8% vs 84.4±5%) in linear actomyosin network simulations. This finding could be reconciled if the force experienced by individual motors in each ensemble is low for slip motors, where they will have maximal bond lifetimes, and low-intermediate for catch bonds, where they may have minimal bond lifetimes. One important distinction, however, is that discrete catch bond ensembles are better crosslinkers than discrete slip bond ensembles, as evidenced by the 1.6-fold increase in total network connectivity in catch bond ensemble simulations compared to those with slip bonds. Not surprisingly, given that both coarse-grained catch and slip bonding ensembles are good binders and can bind maximally two actin-like fibers, this distinction does not hold true for their coarse-grained forms, which have more similar connectivity. In sum, discretized catch bond motor ensembles are useful for their ability to increase network connectivity but are not the most effective force generators, with a slight increase brought about by coarse-graining (Table S3). Increasing evidence that NMMIIB is a good crosslinker and a poor force generator (1, 34) suggests that for modeling these motors, a catch bond method would be suitable and perhaps more realistic than a slip bond method.

### Catch-slip bonding ensembles

Due to the hybrid nature of their force-dependent unbinding dynamics, catch-slip bond motors could be predicted to be the most versatile model. Indeed, in both patch and periodic network simulations, discrete catch-slip ensembles had the highest contractile speeds while also bearing the highest total connectivity. In terms of force generation, contractile patches with catch-slip ensembles bore the highest force generation (~7-10-fold higher than catch or slip ensembles), but in the less permissive periodic linear network simulations, force generation was more comparable (~2-5-fold higher only). Here again, it stands to reason that since force experienced by individual motors may not be very high, catch-slip motors would be the best individual binders of actin-like filaments. Indeed, comparison of individual motor binding shows that catch-slip bond motors are bound 54±5.2% of the time versus 30±5.6% for catch and 16±2.7% for slip bond motors. As with slip and catch bonding ensembles, coarse-graining of catch-slip bond ensembles renders them less effective force generators and drastically reduces total network connectivity (2,900±80 compared to 113,00±6300 for discrete). As such, there is no inherent benefit to coarse-graining catch-slip bonding ensembles. Within a biological context, discrete catch-slip bond ensembles could be the most robust way of simulating non-processive motors that form processive collectives in ensembles. This robustness was shown previously in simulations of gliding filaments on clusters of fixed motors (34), and here we show the same behavior in the context of generic actomyosin contractile systems. Therefore, we suggest that discretized catch-slip bond ensembles provide the best *in silico* approximation for NMMIIA ensembles in the context of contractile systems where connectivity is considered.

### Coarse-graining ensembles

Generally, coarse-graining NMMII ensembles reduces complexity and can speed up computation of simulations by 2 to 3-fold. However, our work shows that coarse-grained ensembles do not generate as much connectivity as discretized ensembles of the same bond type. Comparison among the coarse-grained forms of all bond types revealed insignificant differences between their total connectivity, which averaged around 2,900 connections. This stark contrast to huge differences in connectivity between the discretized versions of each bond type revealed that coarse-graining not only greatly diminishes network connectivity, but also reduces the effect of bond type on connectivity. This is not surprising given the total reduction in motoring fiber binders by 16-fold (from 32 per ensemble to 2 per ensemble). It is important to note that coarse-grained versions of each bond type ensemble still generate vastly different contractile dynamics from one another. This is evident by both contractile rates and force generation estimates for filament patches and for rupture timing and force generation estimates for linear contractile networks. Overall, with the exception of PCM ensembles, which have different emergent properties based on the method of calculating effective bound velocity and binding dynamics, coarse-graining conserves the general relationships between the motor bond types. Catch-slip bond ensembles remain the most effective force-generators (ignoring PCM) and slip ensembles the worst binders. Coarse-graining ensembles can therefore be considered a useful strategy for simulations that seek to describe general principles of contractility in large contractile systems.

### On contractile rings

One long-term application of the work presented herein is to build a predictive model of actomyosin contractility in the cytokinetic contractile ring, where NMMII is the major force generator and also contributes to connectivity (10, 20, 51). Recent *in silico* work on contractile ring dynamics has focused on modeling fission yeast cytokinesis (11, 38–40, 52). Modeling this contractile system is especially powerful due to the availability of published measurements of the absolute abundance of cytoskeletal components in the fission yeast cytokinetic ring (45, 46). Much of this modeling work has used simplified coarse-grained or partially coarse-grained motors to represent NMMII ensembles (39, 40). As such, connectivity of the actomyosin network has largely been simplified. We sought preliminary evidence that the discretized catch and catch-slip motor ensembles we described above could yield contractile ring simulations with realistic network connectivity, even though the model of the ring is generalized and simplified, lacking, for instance, anchorage to a moving plasma membrane-like boundary. Modeling contractile rings bearing each of our different types of motor ensembles revealed similar relationships to those presented above for patches and linear networks (Movie 8; Movie 5). Discretized catch-slip bond ensembles provided the most realistic combination of force generation, contractile speed, and connectivity (Figure 6F-H). Discretized catch bond ensembles were only slightly inferior in terms of speed and force generation, with slip bond and PCM ensembles performing the least optimally. These findings suggest that the motor behaviors presented here will be relevant to diverse actomyosin networks where connectivity is important for force transmission.

**Figure 6.**
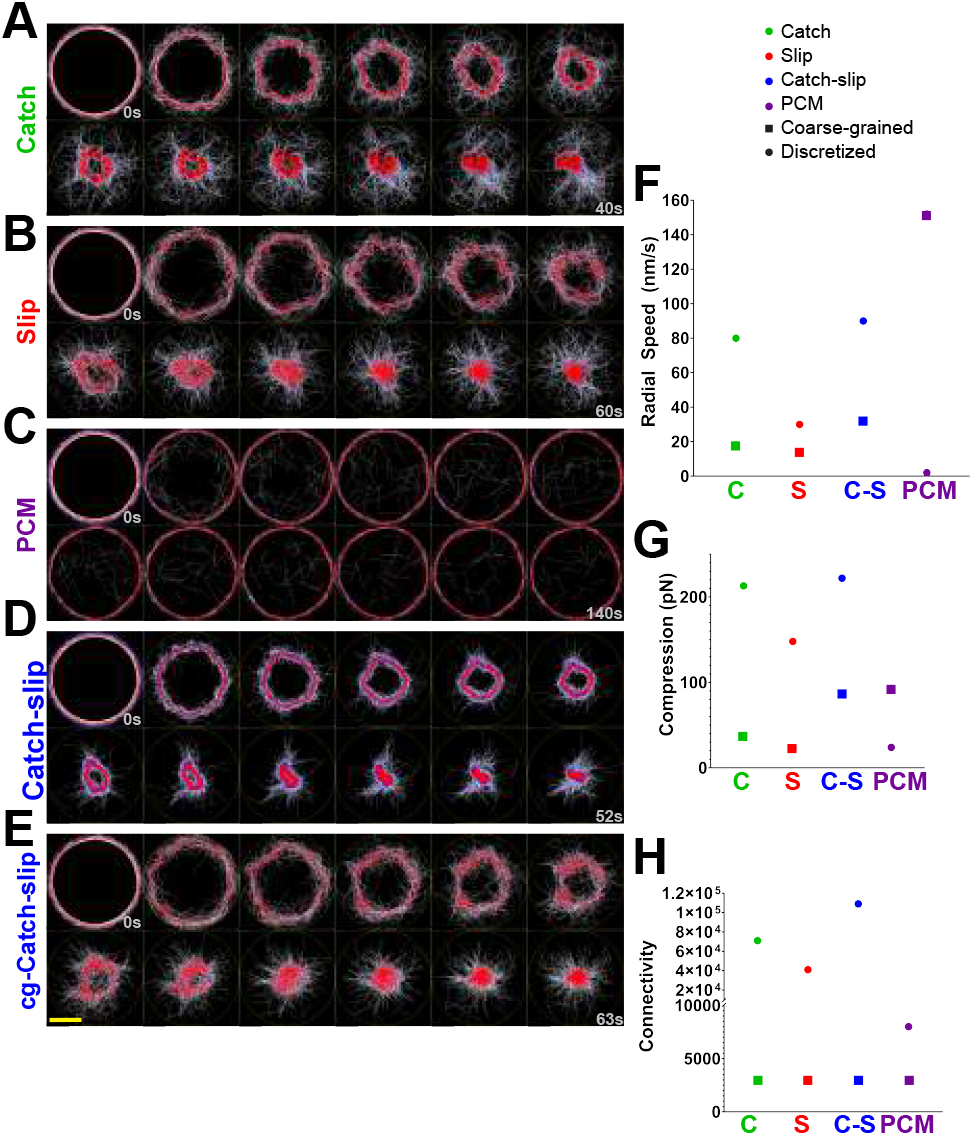
Contractile ring simulations. A-E) Representative timeframes from simulations of fission yeast-sized contractile rings with all four ensemble types (A-D) and coarse-grained catch-slip motor ensembles (E). Scale bar (yellow; bottom left) is 1 µm. F) Maximum radial contractile speed shown for single simulations of rings shown in A-E of each ensemble type with discretized and coarse-grained motors. G) Maximum tension calculated from rings bearing ensembles of each bond type in both discretized and coarse-grained formats. H) Total connectivity of contractile rings bearing ensembles of each bond type with discretizes or coarse-grained motors. Dots in all plots represent data from contractile rings with discretized ensembles; squares represent data from rings with coarse-grained ensembles.

### Perspectives

Going forward, our study of modeling motor ensembles will facilitate more accurate simulations of biological contractile networks. However, several further steps can be taken to increase biological realism beyond the scope of motor ensemble simulations presented here. Motoring activity has been shown to be dispensable for contractility in some systems (17). As such, future work should further focus on separately defining the force contributions of NMMII motoring and crosslinking activity to contractility. Here, we preliminarily demonstrate that motor-dead NMMII ensembles with catch-slip bonds and simulations with no motor ensembles can both still result in partial, though significantly reduced, contractility of network patches (Table S3; Figure S4C-D of the Supporting Material). Actin dynamics are complex and can vary for many reasons. In the work presented here we simulated actin polymerization and depolymerization at both ends of the filament along with a net treadmilling rate. Actin filament buckling can induce severing (13) and depolymerization (53), neither of which we included. Severing can be implemented in Cytosim; bending-dependent depolymerization currently cannot. Future work characterizing the effects of these more complex actin dynamics in contractile systems will undoubtedly provide valuable insight into the mechanisms of contractility. A final, important component of contractile ring dynamics that remains mostly ignored is the contribution of the plasma membrane to the mechanics of contraction. A recent model incorporated fluid membranes (40), but motor ensembles in this study were significantly coarse-grained. Our work in describing the effects of coarse-graining on network connectivity suggests that future work with deformable membranes would further benefit from using discretized motor ensembles.

## AUTHOR CONTRIBUTIONS

D.B.C., A.S.M, and F.N designed the research. D.B.C. carried out most of the simulations and data analysis. M.G. performed some simulations and assisted with data analysis. D.B.C. and A.S.M. wrote the manuscript with support from F.N.

## ACKNOWLEDGEMENTS

The authors thank the members of the Maddox labs, especially Michael Werner, Jenna Perry and Tanner Fadero for critical reading of this manuscript and Shivanandh Kammala for help in data processing. This work was supported by the NIH/NIGMS R01-102390 and NSF 1616661. Simulation work was run on the UNC Chapel Hill Research Computing Longleaf cluster. F.N. is supported by the Gatsby Charitable Foundation.

## SUPPORTING CITATIONS

References (54–59) appear in the Supporting Material.

## FIGURE LEGENDS

**Movie 1. Simulations of force-dependent single motor dynamics.** Simulations for all four motor types are shown in the proximity of a single actin-like filament (white) which is permanently bound to a single Hookean spring (orange). Solid color indicates binding of catch bond (green), slip bond (red), catch-slip bond (blue), or PCM (purple) discrete single motors; brighter coloring indicates bound state. Timesteps are 1 millisecond apart; total simulation time is 0.8 seconds shown at 40 frames/second.

**Movie 2. Ensemble translocation on immobilized filaments.** Four separate simulations of translocating motor ensembles on immobilized filaments each show 40 ensembles, with each ensemble bearing 16 motors of one specified type at one end. Ensemble backbones (yellow lines) appear throughout the simulation, motors (solid colored balls) only appear when they are bound to actin-like filaments (white). Motor ensembles are seeded on the left side and translocate toward the right (barbed) end of actin-like filaments. Top: catch bond motors (green); Top middle: slip motors (red); Bottom middle: catch-slip motors (blue); Bottom: PCM motors (purple). Timesteps are 100 milliseconds apart; total simulation time is 120 seconds shown at 40 frames/second.

**Movie 3. Contractile patches with different crosslinker amounts.** Three simulations of contractile patches (white filaments) within 2D circular spaces (yellow circles). Total number of motor ensembles (red) is the same across all three but crosslinker amount (blue) increases from left to right. Crosslinker amount affects network structure and contractile dynamics. Timesteps are 0.5 seconds apart; total simulation time is 30 seconds shown at 12 frames/second.

**Movie 4. Linear periodic contractile networks.** Simulations of linear networks made of white filaments and blue crosslinkers bearing catch-slip (top) or PCM (bottom) motor ensembles in red. Timesteps are 0.5 seconds apart; total simulation time is 50 seconds shown at 12 frames/second.

**Movie 5. Contractile ring dynamics with discretized catch-slip bond motor ensembles.** Simulation shows a contractile ring formed by seeding filaments (white) in a circular pattern tangential to the simulation space (yellow circle) with crosslinkers (blue) and motors (red). Timesteps are 0.5 seconds apart; total simulation time is 54 seconds shown at 12 frames/second.

**Movie 6. Highly structured and bundled filaments in patch simulations.** Simulation of a contractile patch (as in Movie 3) bearing discrete PCM motor ensembles. These patches are not very contractile and instead form large bundles of mostly linear filaments over time. Only filaments (white) are shown to highlight the bundling of filaments and to show input data for directionality analysis. Timesteps are 0.5 seconds apart; total simulation time is 203 seconds shown at 12 frames/second.

**Movie 7. Ensemble translocation of discrete versus coarse-grained ensembles.** Simulation of discrete (dark blue) or coarse-grained (sky blue) catch-slip ensembles translocating on immobilized actin-like filaments. Dots appear when motors are bound and actively translocating from left to right. Yellow lines represent ensemble rods to which motors are bound. Free parameters for coarse-grained ensembles were fit via parameter sweeps to behave indistinguishably from discretized versions. Timesteps are 100 milliseconds apart; total simulation time is 80 seconds shown at 40 frames/second.

**Movie 8. Contractile ring dynamics with discretized catch bond motor ensembles.** Simulation shows a contractile ring formed by seeding filaments (white) in a circular pattern tangential to the simulation space (yellow circle) with crosslinkers (blue) and motors (red). Timesteps are 0.5 seconds apart; total simulation time is 43 seconds shown at 12 frames/second.

**Table S1.** General Simulation Parameters. Common parameters for elements simulated on contractile patches, linear networks and rings.

| Parameter | Comments | Used for | Ref. |
| --- | --- | --- | --- |
| **ENVIRONMENT** |  |  |  |
| Viscosity = 1 N.s/m<sup>2</sup> | *C. elegans* embryo cytoplasm | ALL | (1) |
| kT = 0.0042 pN.μm | Thermal energy at room temperature | ALL |  |
| Time-step $\tau$ = 1 ms | Movements of the objects are solved every millisecond; simulations are rendered every 0.5 or 1s as indicated in Cytosim configuration files. | ALL | |
| **SIMULATION SPACE** |  |  |  |
| semi-periodic | Space is periodic in the X direction (length 2μm) and bounded in the Y direction (width 0.2 μm) | MM1 |  |
| semi-periodic | Space is periodic in the X direction (length 12μm) and bounded in the Y direction (width 4 μm) | MM2 |  |
| disc | Space is a disc with a diameter of 3 μm | MM3 |  |
| semi-periodic | Space is periodic in the X direction (length 9.424 μm) and bounded in the Y direction (width 1 μm) | MM4 |  |
| disc | Space is a disc with a diameter of 3 μm | MM5 |  |
| **ACTIN-LIKE FILAMENTS** |  |  |  |
| Quantity = 360 | This value is extrapolated using *S. pombe* measurements | MM3-MM5 | (2, 3) |
| Length:<br>2 μm<br>12.01 μm<br>1 ± 0.3 μm | Allows length distribution around 1 μm | MM1<br>MM2<br>MM3-MM5 | (2) |
| Rigidity:<br>20 pN.μm<sup>2</sup><br>0.06 pN.μm<sup>2</sup> |  | MM2<br>All other | (5) |
| **Plus (barbed) ends:**<br>growth speed = 0.1 μm/s<br>shrinking speed = 0.004 μm/s |  | MM3-MM5 | (5) |
| **Minus (pointed) ends:**<br>growth speed = 0.002 μm/s<br>shrinking speed = 0.1 μm/s |  | MM3-MM5 | (5) |
| **MOTOR SUBUNITS** |  |  |  |
| **All non-PCM Motors** |  |  |  |
| Stiffness = 100 pN/μm | Empirical; 200 pN/μm for coarse-grained motor ensembles in all simulations | ALL |  |
| Unbinding force = 5 pN |  | ALL | (8) |
| **Discrete Catch** |  |  |  |
| Binding rate = 0.2 /s |  | ALL | (6) |
| Binding range = 0.06 μm | Estimated from electron micrographs of *in vitro* reconstituted NMM2 | ALL | (7) |
| Unbinding rate = 1.71/s |  | ALL | (6) |
| Force-dependent unbinding | Catch equation; Eq. 6 | ALL |  |
| Max speed ($V_0$) = 0.15 μm/s | | ALL | (12, 13) |
| Stall force ($F_s$) = 3.85 pN | | ALL | (8, 14) |
| $\alpha_{\text{catch}} = 0.92$ | Weighted prefactor as described in Stam *et al.* | ALL | (10, 16) |
| $\chi_{\text{catch}} = 0.607681$ pN | Derived from catch-slip equation (bond length over kT) | ALL | (16) |
| **Coarse Grained Catch** |  |  |  |
| Binding rate = 2.48 /s | Determined by a parameter sweep (see Figure S4B) | MM2-MM6 |  |
| Binding range = 0.12 μm | Double the value for discrete version of motor | MM2-MM6 | (7) |
| Unbinding rate = 0.9 /s | Determined by a parameter sweep (see Figure S4B) | MM2-MM6 |  |
| Force-dependent unbinding | Catch equation (Eq. 6) | MM2-MM6 |  |
| Max speed ($V_0$) = 0.145 $\mu\text{m/s}$ | Determined by a parameter sweep (see Figure S4B) | MM2-MM6 | |
| Stall force ($F_s$) = 4 pN | Determined by a parameter sweep (see Figure S4B) | MM2-MM6 | |
| $\alpha_{\text{catch}} = 0.92$ | Weighted prefactor as described in Stam *et al.* | MM2-MM6 | (10, 6) |
| $\chi_{\text{catch}} = 0.607681$ pN | Derived from catch-slip equation (bond length over $kT$) | MM2-MM6 | (10) |
| **Discrete Slip** |  |  |  |
| Binding rate = 0.2 /s |  | ALL | (6) |
| Binding range = 0.06 $\mu\text{m}$ | Estimated from electron micrographs of *in vitro* reconstituted NMM2 | ALL | (7) |
| Unbinding rate = 1.71/s |  | ALL | (6) |
| Force-dependent unbinding | Kramers' theory (Eq. 2) | ALL |  |
| Max speed ($V_0$) = 0.15 $\mu\text{m/s}$ | | ALL | (12, 13) |
| Stall force ($F_s$) = 3.85 pN | | ALL | (8, 14) |
| **Coarse Grained Slip** |  |  |  |
| Binding rate = 1.32 /s | Determined by a parameter sweep (see Figure S4B) |  |  |
| Binding range = 0.12 $\mu\text{m}$ | Double the value for discrete version of motor | MM2-MM6 | (7) |
| Unbinding rate = 0.8 /s | Determined by a parameter sweep (see Figure S4B) | MM2-MM6 |  |
| Force-dependent unbinding | Kramers' theory; Eq. 2 | MM2-MM6 |  |
| Max speed ($V_0$) = 0.137 $\mu\text{m/s}$ | Determined by a parameter sweep (see Figure S4B) | MM2-MM6 | |
| Stall force ($F_s$) = 15 pN | Determined by a parameter sweep (see Figure S4B) | MM2-MM6 | |
| **Discrete Catchslip** |  |  |  |
| Binding rate = 0.2 /s |  | ALL | (6) |
| Binding range = 0.06 $\mu\text{m}$ | Estimated from electron micrographs of *in vitro* reconstituted NMM2 | ALL | (7) |
| Unbinding rate = 1.71/s |  | ALL | (6) |
| Force-dependent unbinding | Catchslip Equation; Eq. 5 | ALL | (10) |
| Max speed ($V_0$) = 0.15 $\mu\text{m/s}$ | | ALL | (12, 13) |
| Stall force ($F_s$) = 3.85 pN | | ALL | (8, 14) |
| $\alpha_{\text{catch}} = 0.92$ | Weighted prefactor as described in Stam *et al.* | ALL | (10, 6) |
| $\alpha_{\text{slip}} = 0.08$ | Weighted prefactor as described in Stam *et al.* | ALL | (10, 6) |
| $\chi_{\text{catch}} = 0.607681$ pN | Derived from catch-slip equation (bond length over $kT$) | ALL | (10) |
| $\chi_{\text{slip}} = 0.097229$ pN | Derived from catch-slip equation (bond length over $kT$) | ALL | (10) |
| **Coarse Grained Catchslip** |  |  |  |
| Binding rate = 3.6 /s | Determined by a parameter sweep (see Figure S4B) | MM2-MM6 |  |
| Binding range = 0.12 $\mu\text{m}$ | Double the value for discrete version of motor | MM2-MM6 | |
| Unbinding rate = 0.8 /s | Determined by a parameter sweep (see Figure S4B) | MM2-MM6 |  |
| Force-dependent unbinding | Catch-slip Equation (Eq. 5) | MM2-MM6 | (10) |
| Max speed ($V_0$) = 0.146 $\mu\text{m/s}$ | Determined by a parameter sweep (see Figure S4B) | MM2-MM6 | |
| Stall force ($F_s$) = 5 pN | Determined by a parameter sweep (see Figure S4B) | MM2-MM6 | |
| $\alpha_{\text{catch}} = 0.92$ | Weighted prefactor as described in Stam *et al.* | MM2-MM6 | (10, 6) |
| $\alpha_{\text{slip}} = 0.08$ | Weighted prefactor as described in Stam *et al.* | MM2-MM6 | (10, 6) |
| $\chi_{\text{catch}} = 0.607681$ pN | Derived from catch-slip equation (bond length over $kT$) | MM2-MM6 | (10) |
| $\chi_{\text{slip}} = 0.097229$ pN | Derived from catch-slip equation (bond length over $kT$) | MM2-MM6 | (10) |
| **PCM Motors** |  |  |  |
| Binding rate = 0.2 /s | Listed as 'k_01' in PCM configuration files | ALL | (6) |
| Binding range = 0.06 $\mu\text{m}$ | Estimated from electron micrographs of *in vitro* reconstituted NMM2 | ALL | (7) |
| Unbinding rate = 1.71 /s | Listed as 'k_20' in PCM configuration files | ALL | (6) |
| Unbinding relationship | PCM method; Eq. 3 | ALL | (11) |
| Max speed ($V_0$) = 0.6 $\mu\text{m/s}$ | Estimated from Erdmann model | ALL | (11) |
| Stall force = 12.3 pN | Estimated from Erdmann model | ALL | (11) |
| Force-speed relationship | PCM method; Eq. 4 | ALL | (11) |
| **Motor Ensembles** |  |  |  |
| Quantity = 360 | This value is extrapolated using *S. pombe* measurements | MM3-MM5 | (2, 3) |
| Valence = 32 subunits/ensemble | Simulated as 2 actin-binding agents of 16 motors in aggregate method) | MM3-MM5 | (12) |
| Length = 0.3 $\mu\text{m}$ | Estimated from electron micrographs of reconstituted NMM2 | MM3-MM5 | (7, 12, 16-18) |
| Rigidity = 100 pN.$\mu\text{m}^2$ | | MM3-MM5 | |
| Diffusion | Ensembles are simulated as a line segment with motors attached at ends. As such their diffusion is calculated by the over-dampened Langevin equation in Cytosim | MM3-MM5 |  |
| **CROSSLINKERS** |  |  |  |
| Quantity = [1,000 - 12,000] | Performed parameter sweeps to establish optimal amount | MM3-MM5 |  |
| Length = 0.04 $\mu\text{m}$ | Approximated from anillin size | MM3-MM5 | |
| Binding rate = 10/s |  | MM3-MM5 |  |
| Binding range = 0.050 $\mu\text{m}$ | | MM3-MM5 | (15) |
| Unbinding rate:<br>0.3 /s for myosin-binders<br>0.1 /s for actin-binders |  | MM3-MM5 |  |
| Unbinding force = 5 pN |  | MM3-MM5 |  |
| Stiffness = 100 pN/$\mu\text{m}$ | | MM3-MM5 | |

**Table S2.** Ensemble Translocation Simulations. Translocation speed (nm/s) and binding percentage data for all ensemble types on immobilized actin-like filaments. Numbers reported are mean ± standard deviation. Sample size is 200 ensembles for each.

| Ensemble | Translocation speed (nm/s) | % Bound (ensembles) |
| --- | --- | --- |
| **Catch** | 130±3.84 | 76±4.85 |
| **cg-Catch** | 130±2.85 | 76±2.4 |
| **Slip** | 121±6.39 | 59±4.74 |
| **cg-Slip** | 121±5.04 | 59±3.87 |
| **Catch-slip** | 135±4.17 | 87±2.88 |
| **cg-Catch-slip** | 135±2.48 | 87±1.46 |
| **PCM** | 51±11 | 100±0.41 |
| **cg-PCM** | 299±0.61 | 98±0.7 |

**Table S3.**
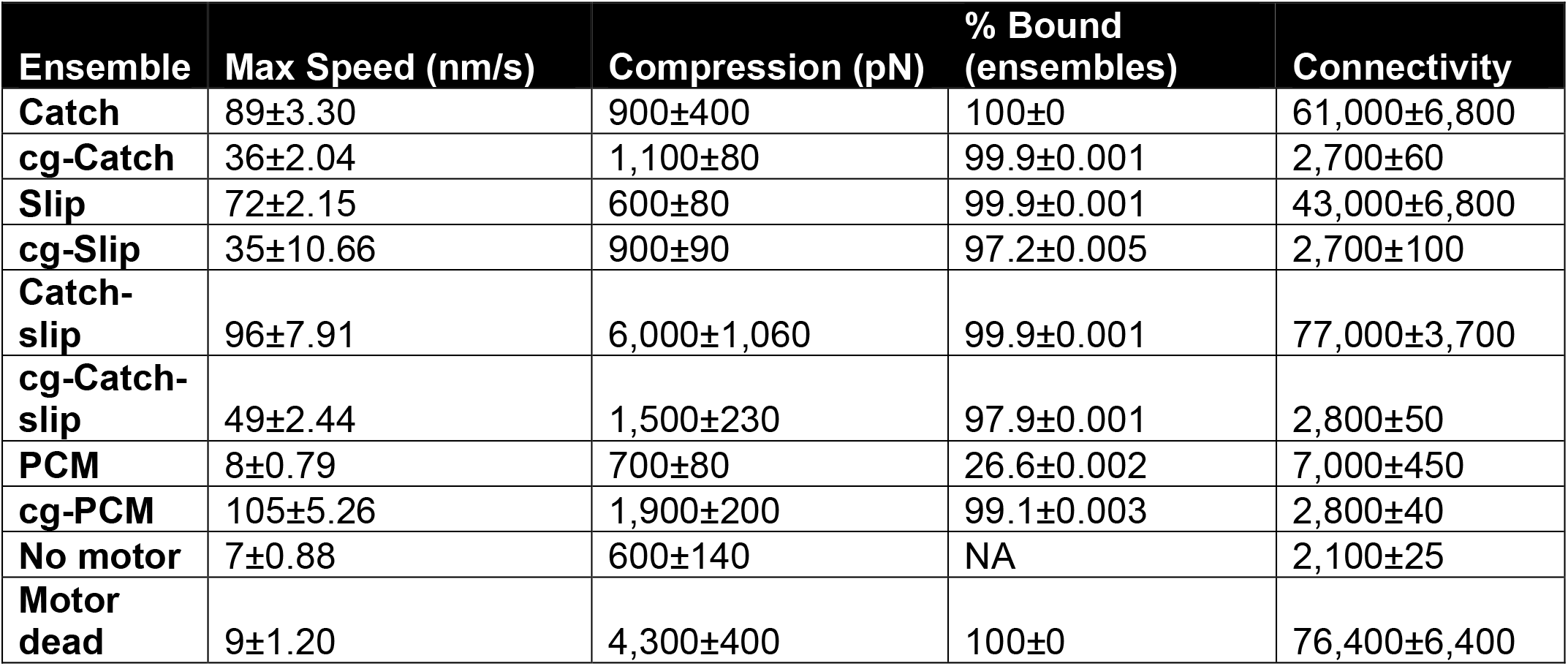
Contractile Patch Simulations. Maximum radial contractile speed (nm/s), network tension (pN) and connectivity for all ensemble types on contractile patches. Numbers reported are mean ± standard deviation. Sample size is 10 independent simulations for each.

| **Ensemble** | **Max Speed (nm/s)** | **Compression (pN)** | **% Bound (ensembles)** | **Connectivity** |
| --- | --- | --- | --- | --- |
| **Catch** | 89±3.30 | 900±400 | 100±0 | 61,000±6,800 |
| **cg-Catch** | 36±2.04 | 1,100±80 | 99.9±0.001 | 2,700±60 |
| **Slip** | 72±2.15 | 600±80 | 99.9±0.001 | 43,000±6,800 |
| **cg-Slip** | 35±10.66 | 900±90 | 97.2±0.005 | 2,700±100 |
| **Catch-slip** | 96±7.91 | 6,000±1,060 | 99.9±0.001 | 77,000±3,700 |
| **cg-Catch-slip** | 49±2.44 | 1,500±230 | 97.9±0.001 | 2,800±50 |
| **PCM** | 8±0.79 | 700±80 | 26.6±0.002 | 7,000±450 |
| **cg-PCM** | 105±5.26 | 1,900±200 | 99.1±0.003 | 2,800±40 |
| **No motor** | 7±0.88 | 600±140 | NA | 2,100±25 |
| **Motor dead** | 9±1.20 | 4,300±400 | 100±0 | 76,400±6,400 |

**Table S4.** Linear Periodic Network Simulations. Maximum tension (pN), binding percentages, and connectivity reported for all ensemble types on linear periodic contractile networks. Numbers reported are mean ± standard deviation. Sample size is 20 independent simulations for each.

| **Ensemble** | **Peak tension (pre fail) (pN)** | **Peak compression (post fail) (pN)** | **% Bound (ensembles)** | **Connectivity** |
| --- | --- | --- | --- | --- |
| **Catch** | 21,000±1160 | 2,400±930 | 84.0±8.0 | 73,000±8,400 |
| **cg-Catch** | 4,000±430 | 420±280 | 98.2±1.2 | 2,900±60 |
| **Slip** | 7,800±560 | 870±150 | 84.4±5.0 | 44,000±5,000 |
| **cg-Slip** | 3,500±300 | 660±140 | 94.6±1.1 | 2,900±60 |
| **Catch-slip** | 39,200±2,100 | 8,600±2,700 | 94.8±6.5 | 113,000±6,400 |
| **cg-Catch-slip** | 5,300±470 | 720±230 | 98.9±1.5 | 2,900±80 |
| **PCM** | 1,300±1,100 | 60±300 | 28.8±0.2 | 8,000±700 |
| **cg-PCM** | 10,300±8,700 | 1,000±230 | 98.1±0.4 | 2,800±30 |

**Figure S1.**
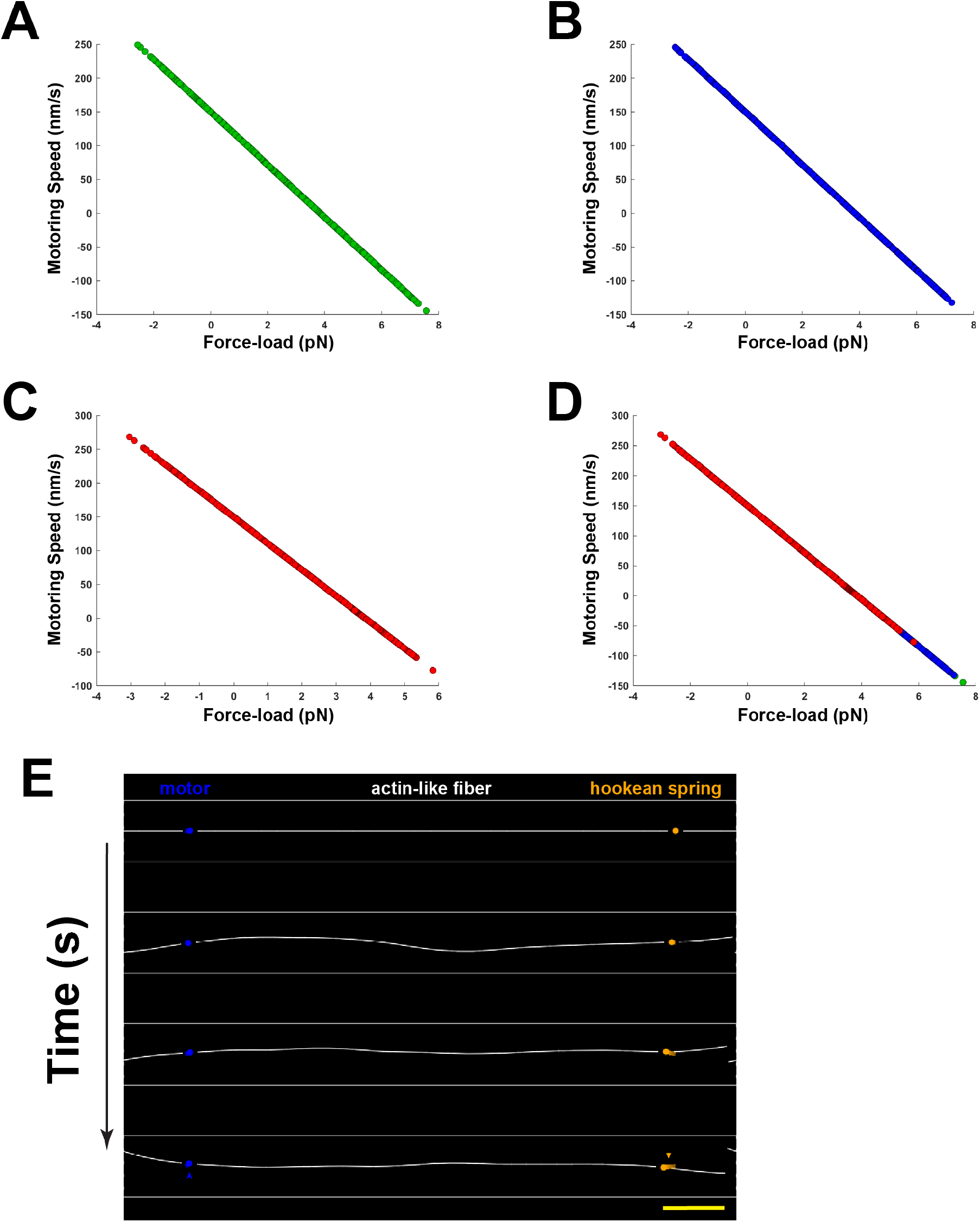
Single motor force-dependent velocity. A-C) The force-velocity relationship for all three (not PCM) bond types follows Eq. 10, where load and velocity are linearly related. D) Overlaid data from A-C showing they all follow the same relationship in Cytosim. Each curve is built from -100,000 individual data points extracted from running simulations. E) Simulation timeframes from a laser trap simulation showing a motor (blue) and a Hookean spring (orange) both bound to an actin-like filament (white). As the simulation progresses the motor pulls on the fiber which causes force to be exerted on the motor by the Hookean spring holding on to the filament. Displacement of motor and spring is visualized by the ‘stretching’ of both components (arrowheads). Scale bar represents 100nm.

**Figure S2.**
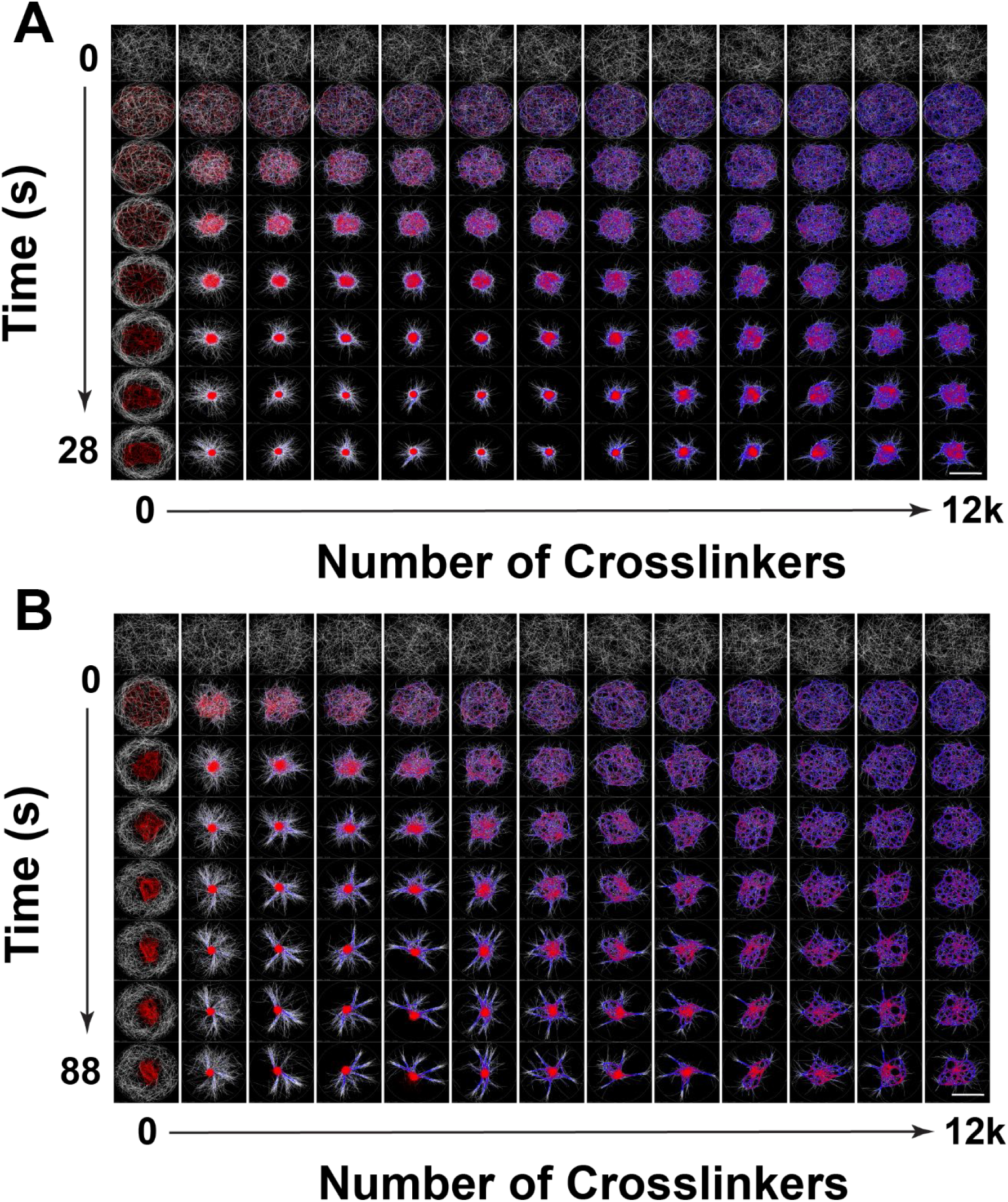
Contractile patch simulation comparison between discrete and coarse-grained ensembles. A-8) Representative simulations of contractile patches. Actin-like filaments (white) are seeded with discrete (A) or coarse-grained (8) motor ensembles (red) and varying amount of crossl inkers (blue) ranging from 0 to 12,000 (along the x-axis). Simulation snapshots are shown from top (Os) to bottom (24s) with 4s intervals for A and from top (Os) to bottom (88s) with 11 second intervals for 8. Scale bar (bottom right) is 1.5μm.

**Figure S3.**
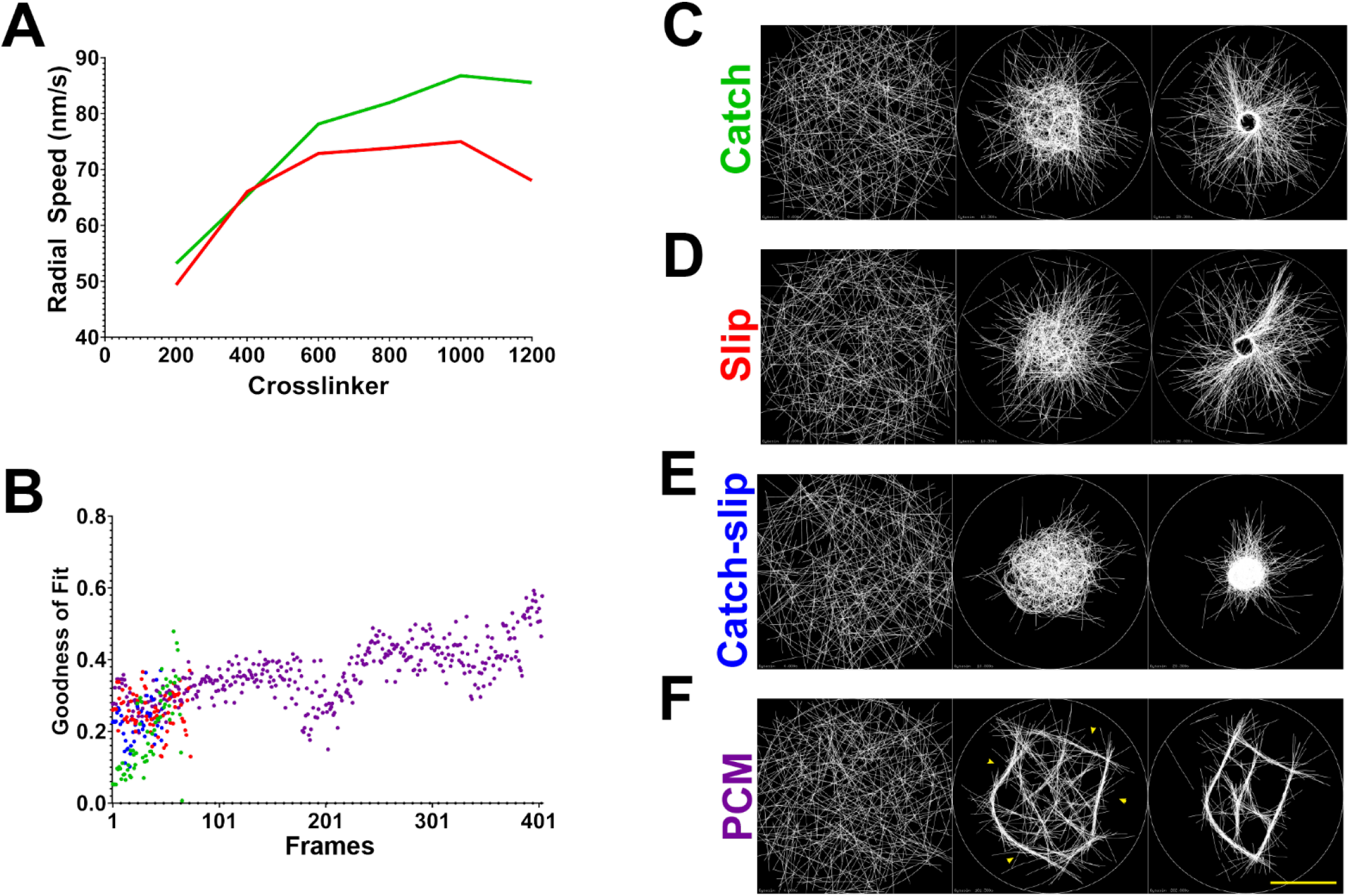
Contractile patch simulation analysis. A) Finer crosslinker sweep as performed for all ensemble types, showing maximum contractile radial speed here from 200 to 1200 crosslinkers (in intervals of 200). Peaks for both catch (green) and slip (red) discretized ensemble types appear at 1000 crosslinkers. 8) Directonality (order) analysis of actin-like filaments (shown in C-F) for each discrete motor ensemble type. Each dot represents the average of 5 separate simulations all ran at peak connectivity for each ensemble type taken every 0.5 seconds (401 frames; total 200.5 seconds). All but PCM (purple) simulations finished contracting before 50s (100 frames). C-F) Representative starting, middle and late timeframes from patch simulations for each discretized ensemble type. White lines are actin-like filaments; white faint circle is the simulation space. Large groups of bundled filaments are obvious in middle timepoints only for PCM simulations (arrowheads). Scale bar (bottom right) is 11μm.

**Figure S4.**
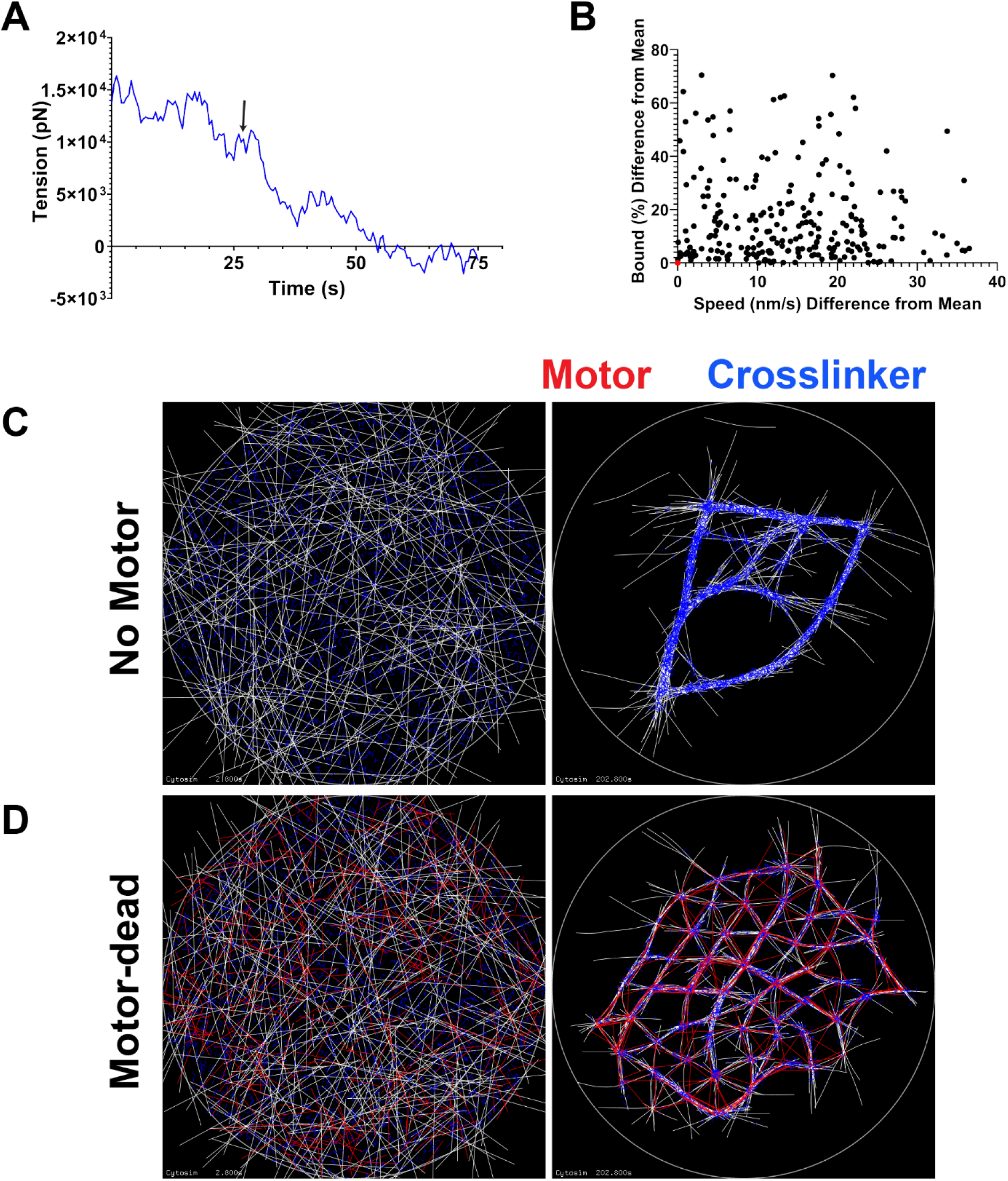
Supplemental data. A) Representative network tension for linear contractile networks over time. Before rupture (black arrow) the tension is large and expansile (positive values on the graph), after rupture tension drops and becomes contractile as filaments contract into local asters. B) Representative graph of coarse-graining parameter sweeps in ensemble translocation simulations. Graph shows deviation from the mean of discrete ensembles for binding percentage (Y-axis) and translocation speed (X-axis). Each dot represents the outputs for a single simulation (average of 40 ensembles per simulation) where binding parameters and motoring parameters were altered. The single red dot near the origin represents a simulation whose swept parameters resulted in outputs near-indistinguishable from the discrete version of the same ensemble. C) Representative simulation of contractile patches with crosslinkers (blue) but no motor ensembles showing partial contraction after 200s (right frame). 0) Representative simulation of contractile patches with crosslinkers (blue) and motor-dead ensembles (red) showing partial contraction after 200s (right frame).

---

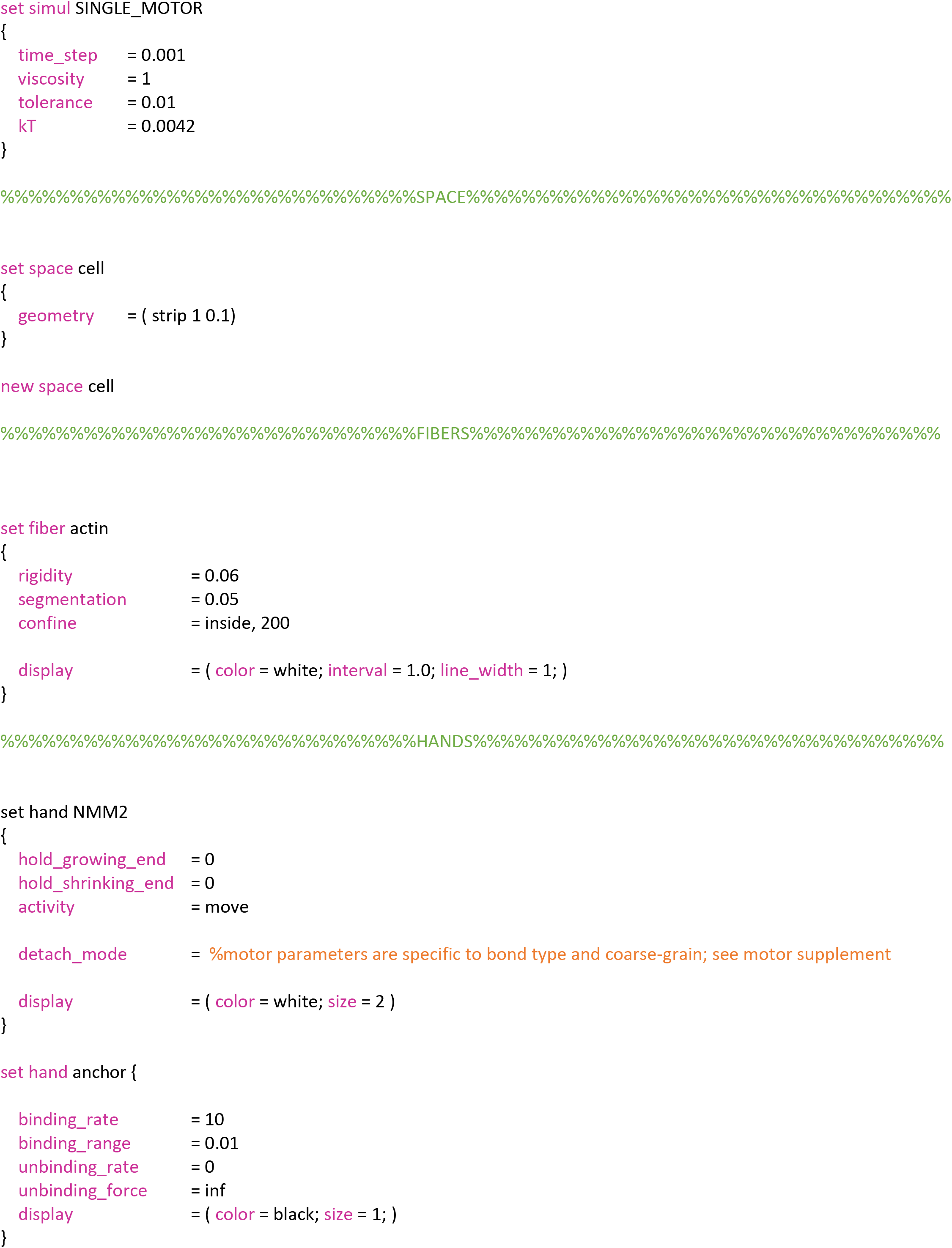
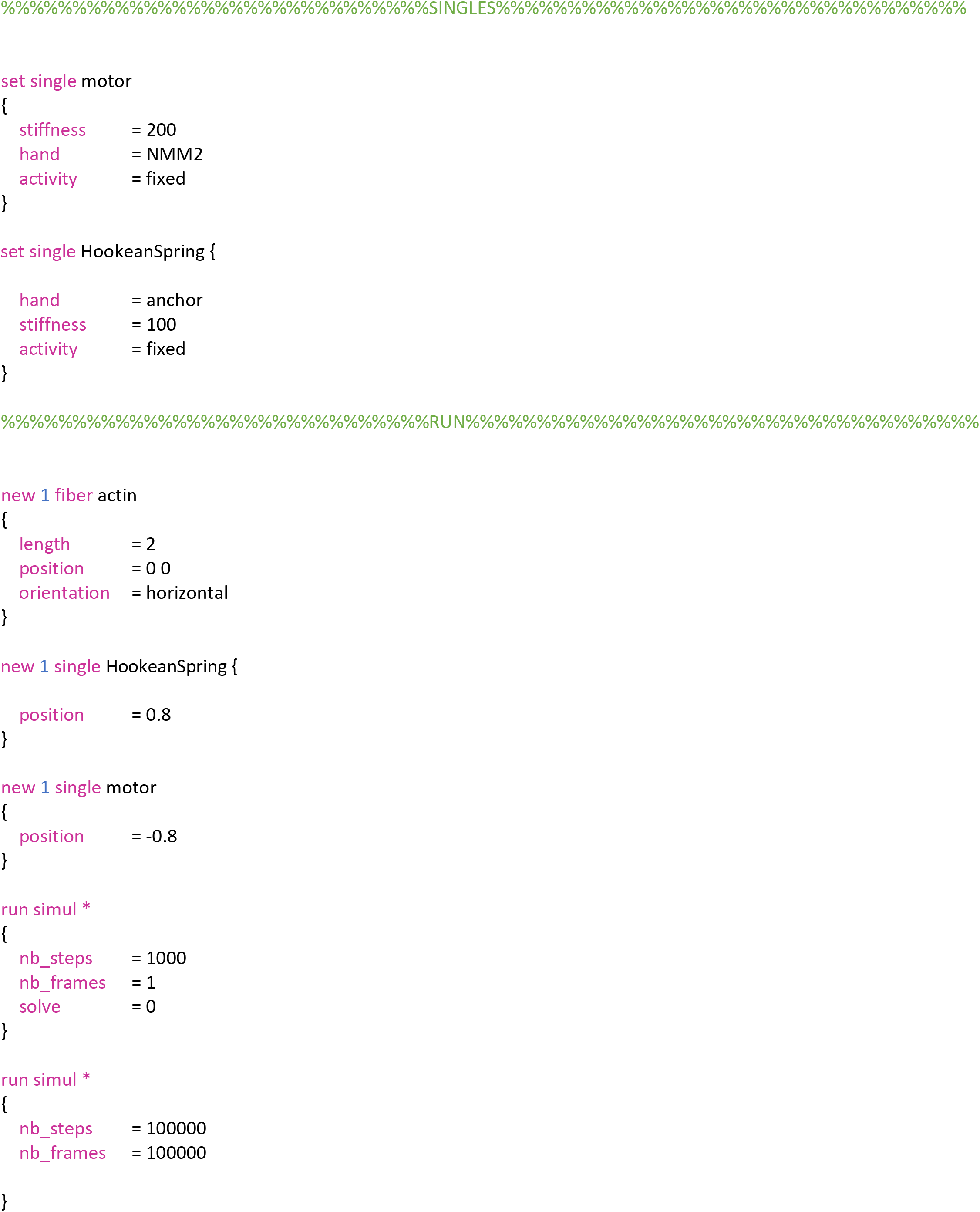
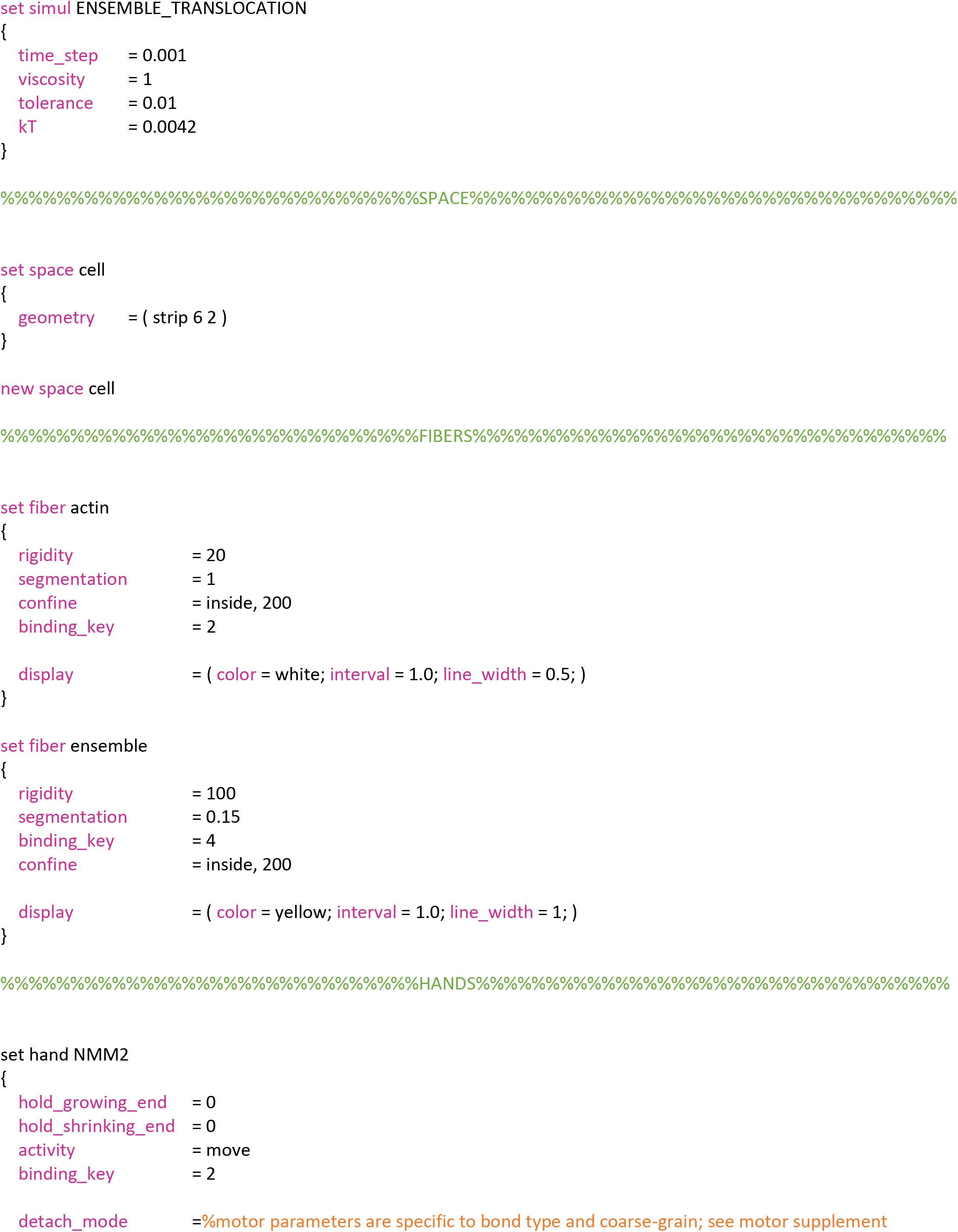
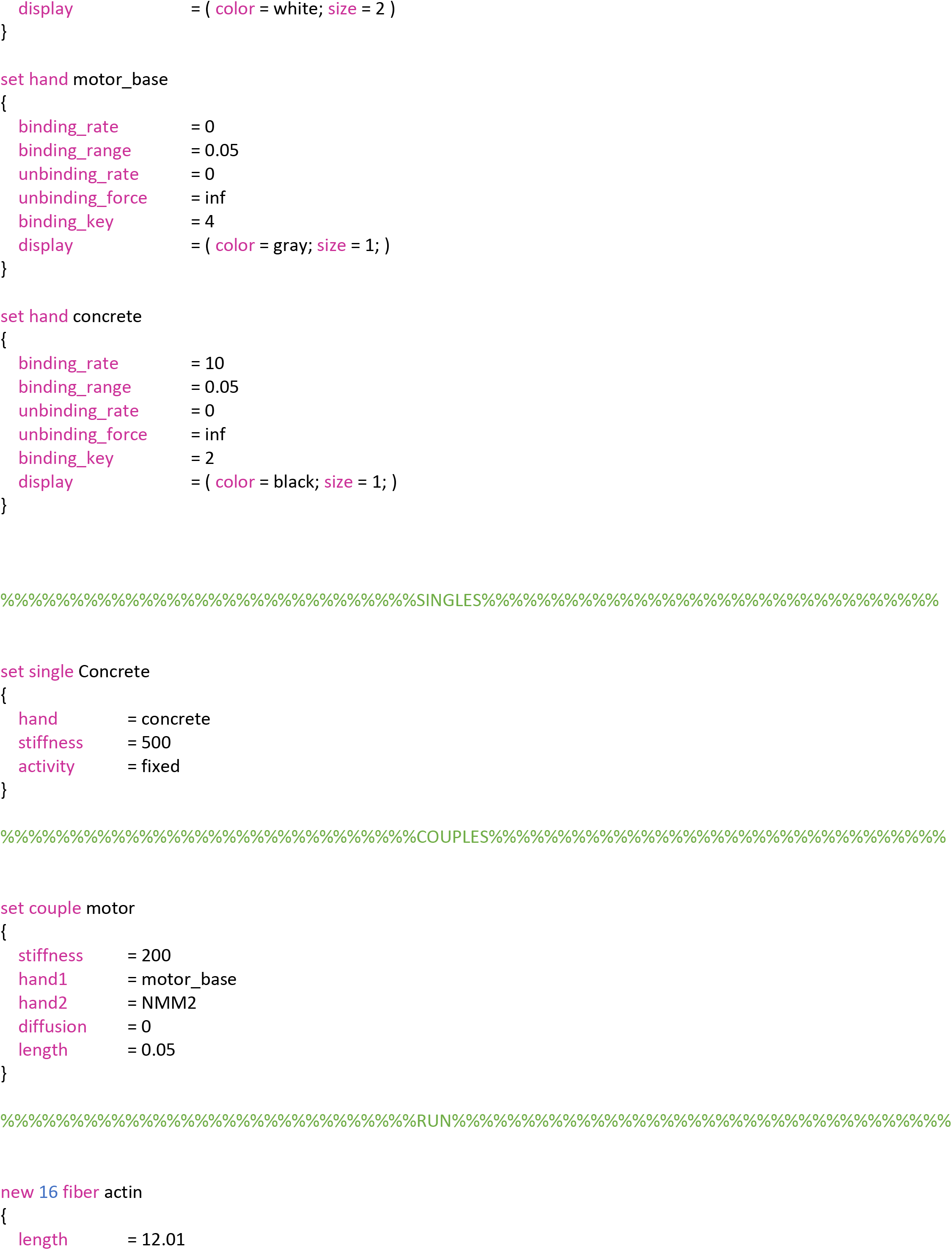
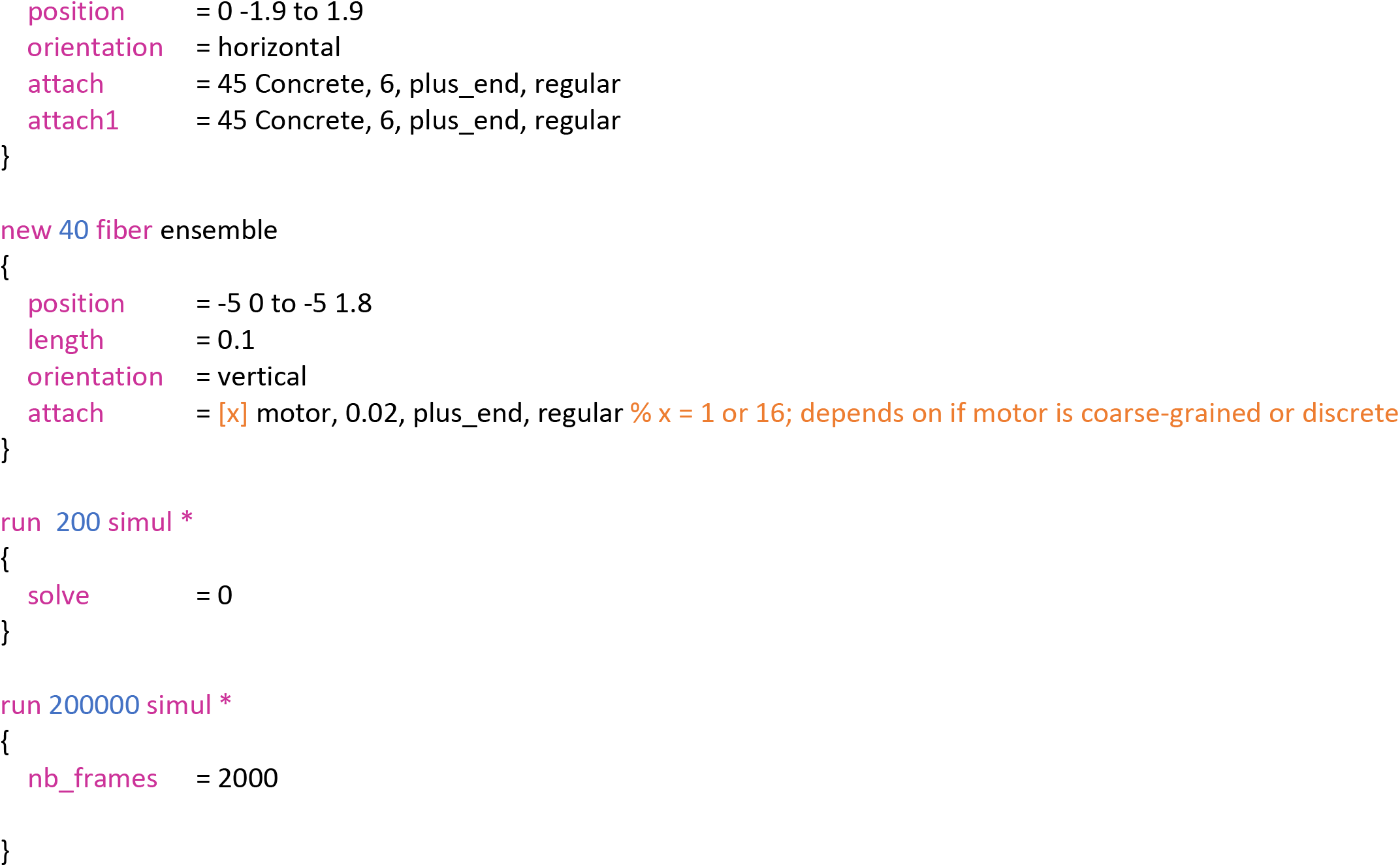
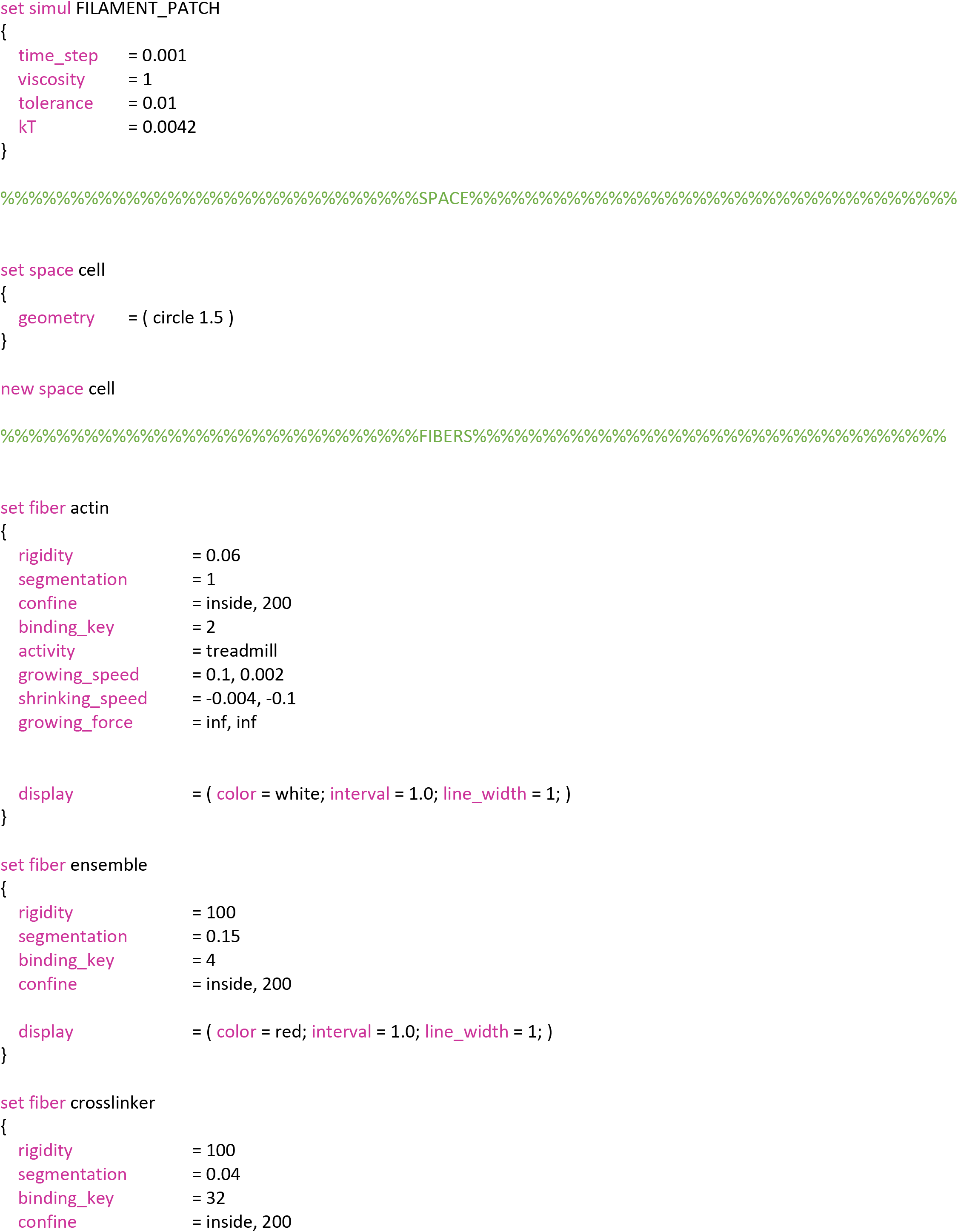
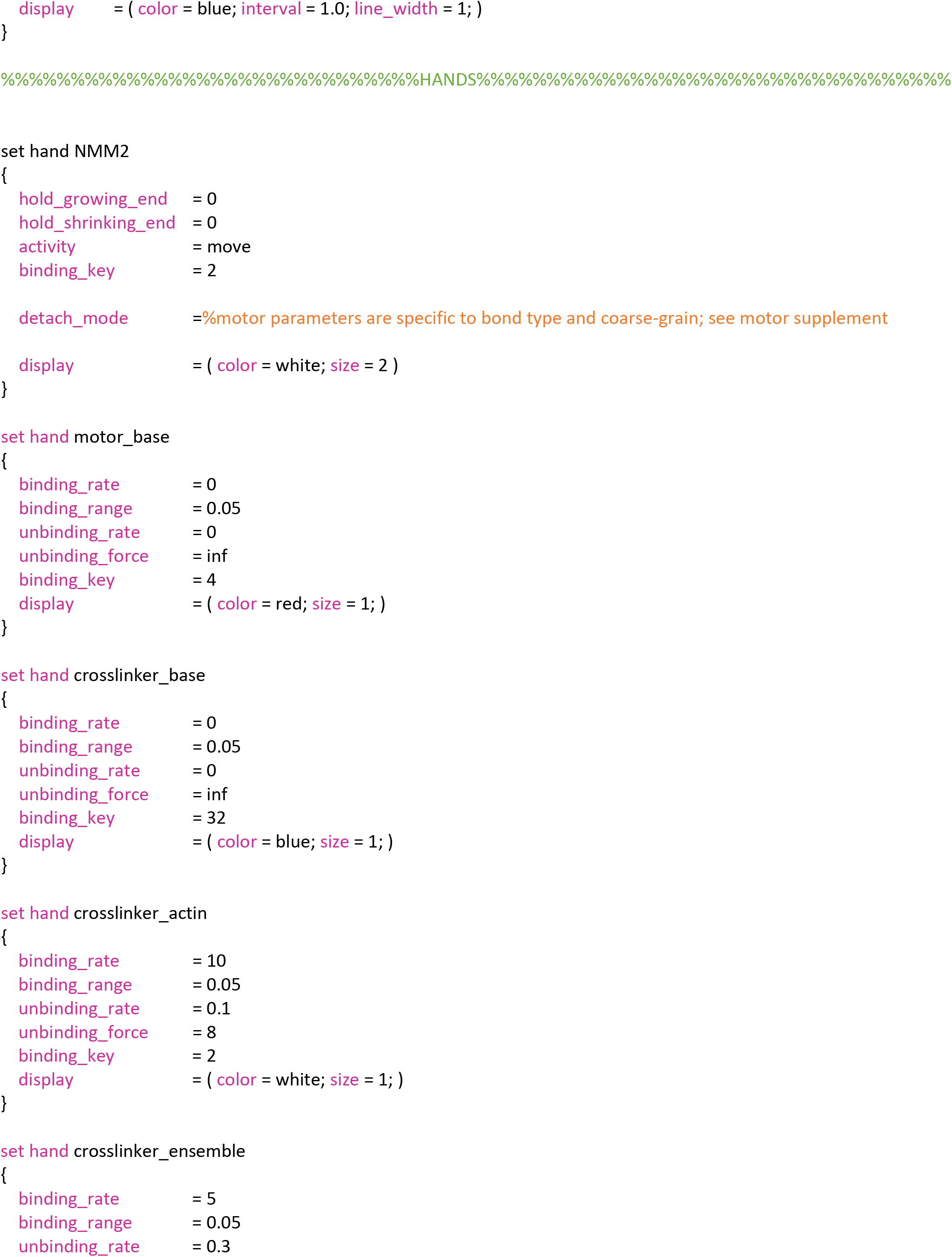
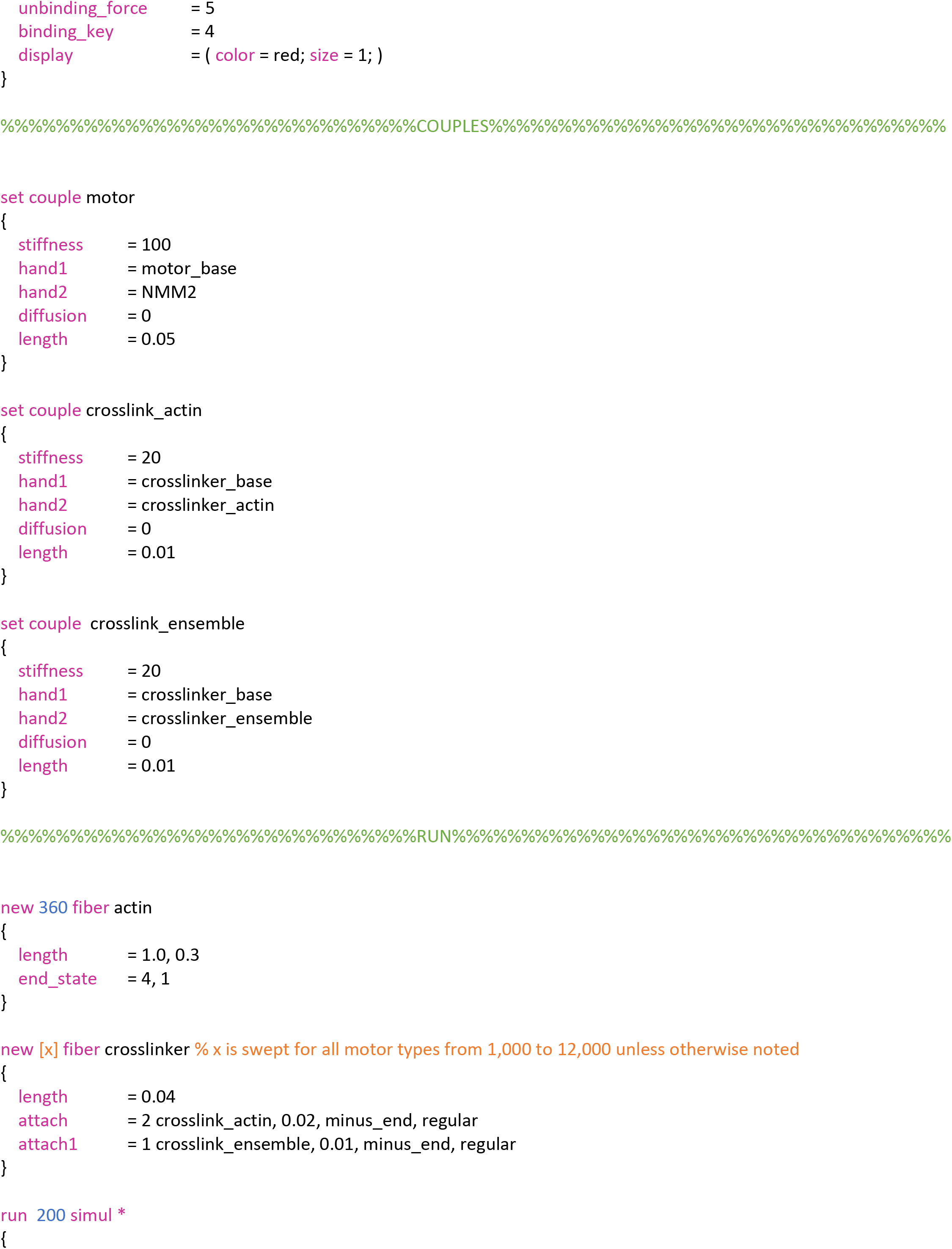
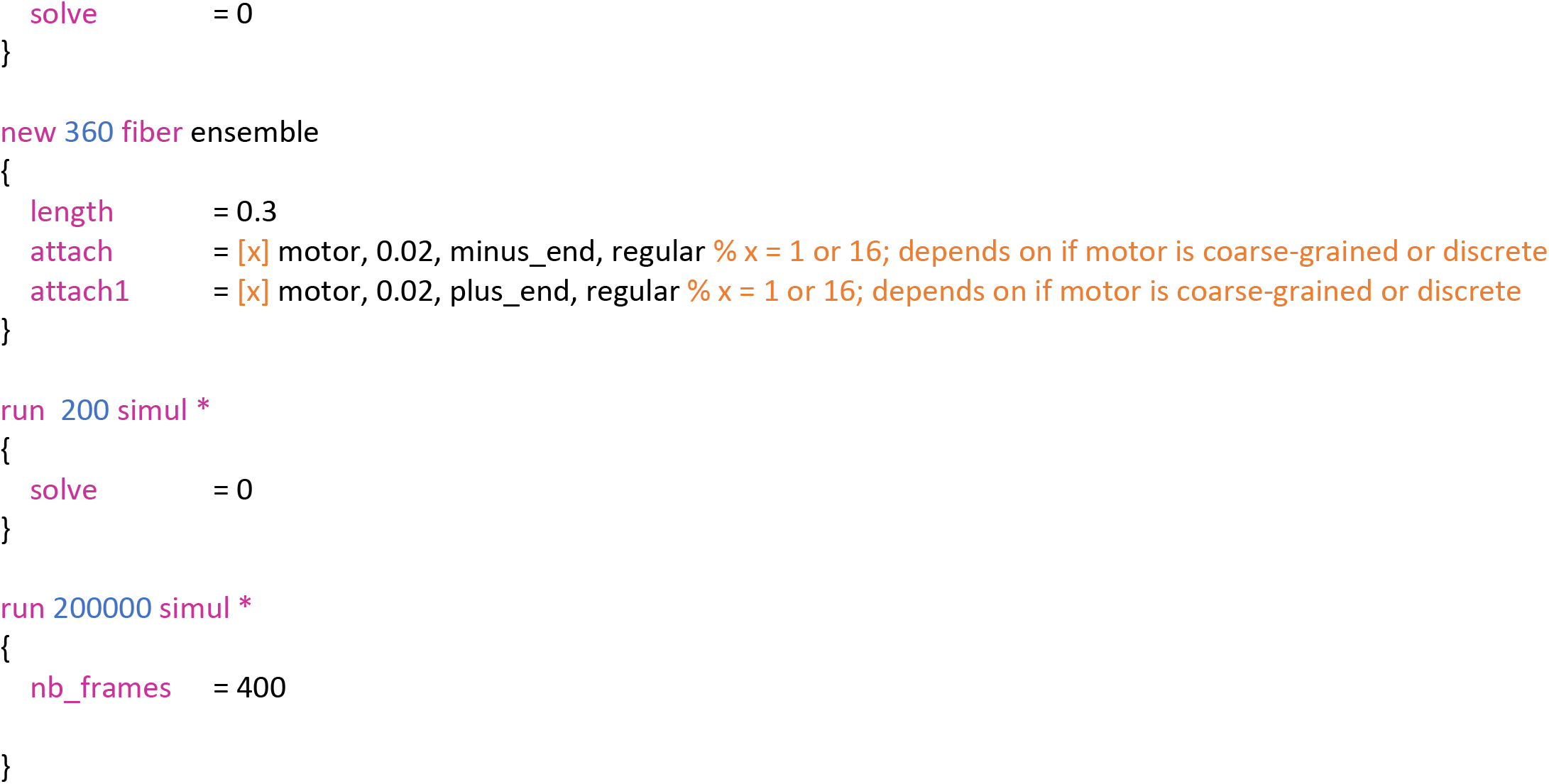
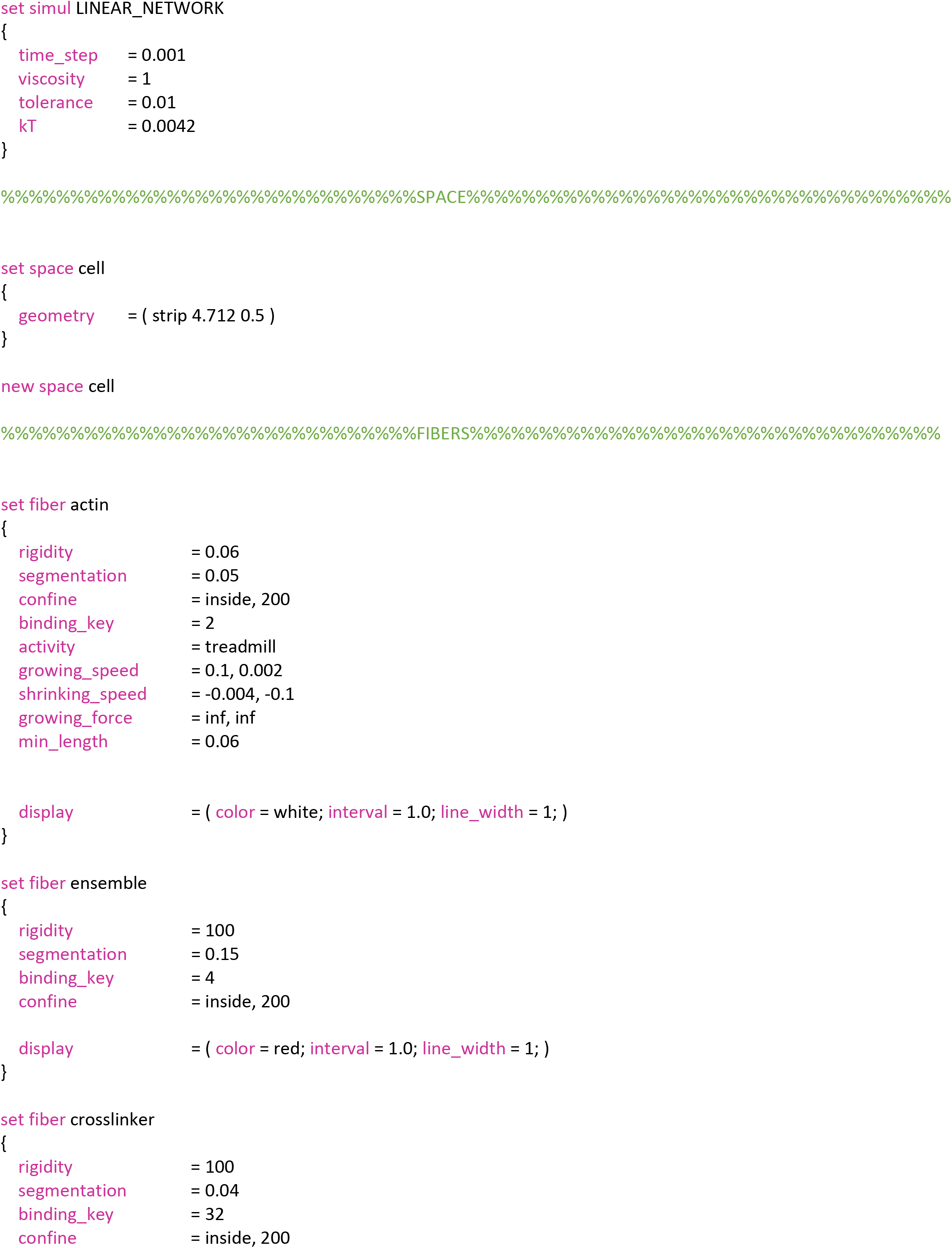
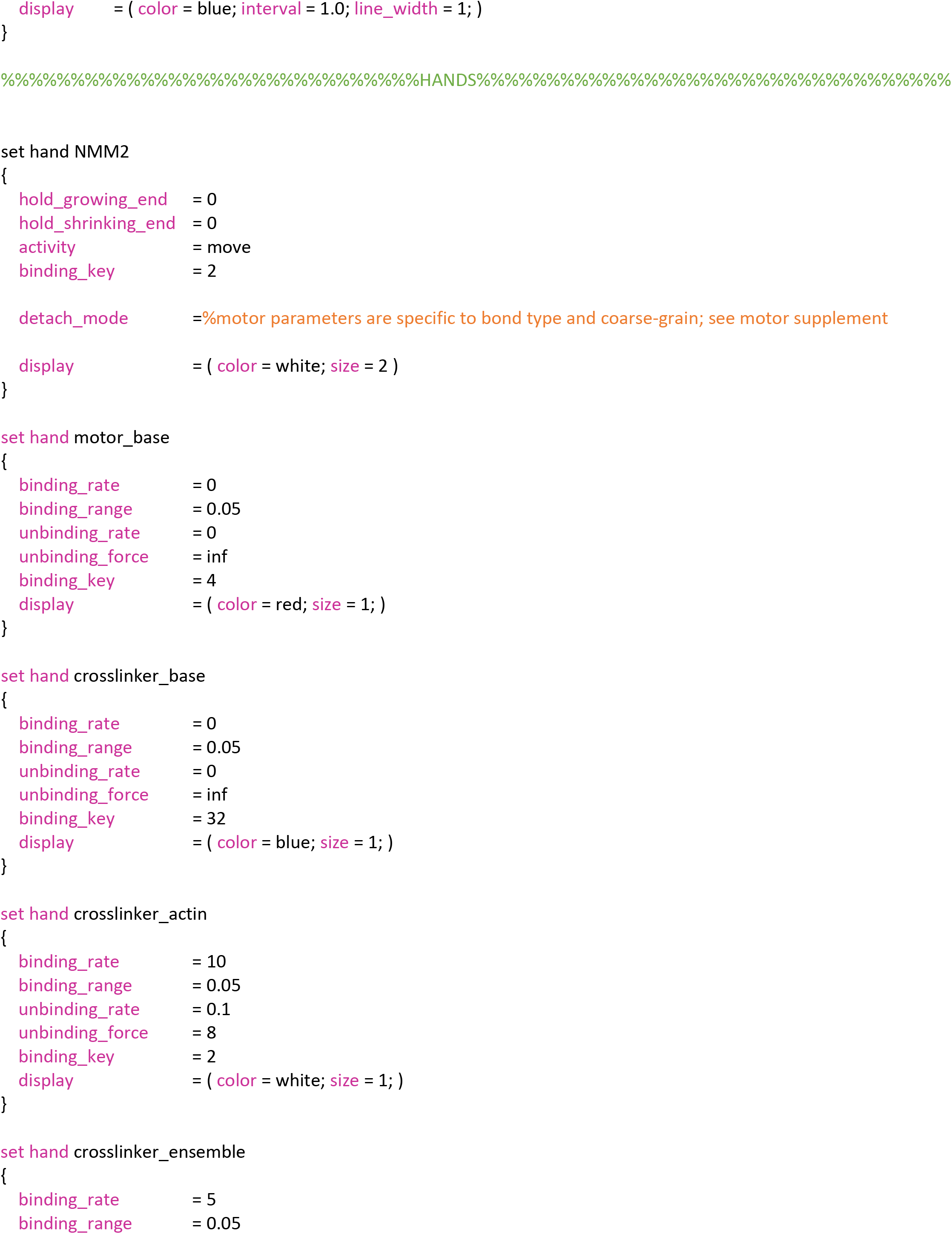
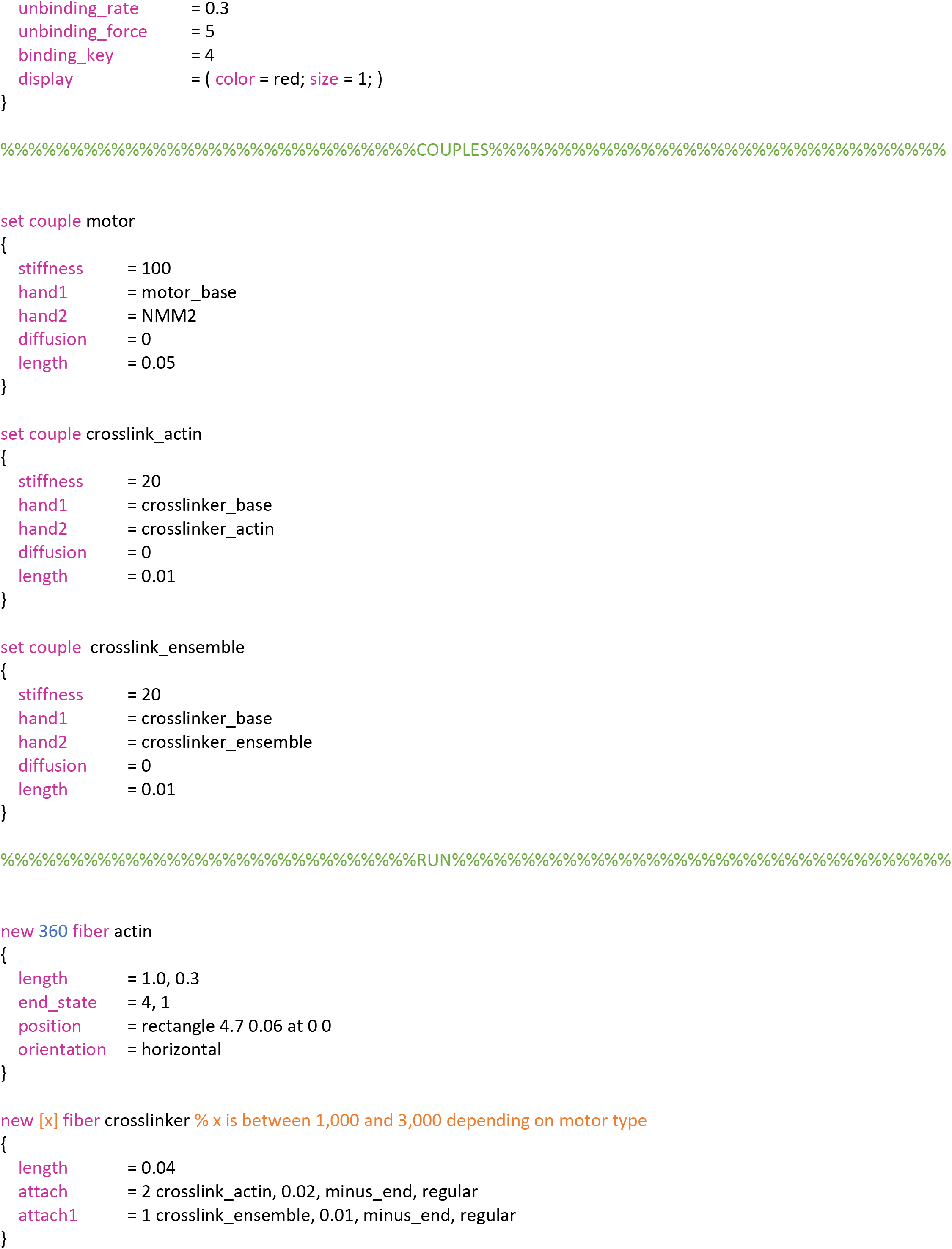
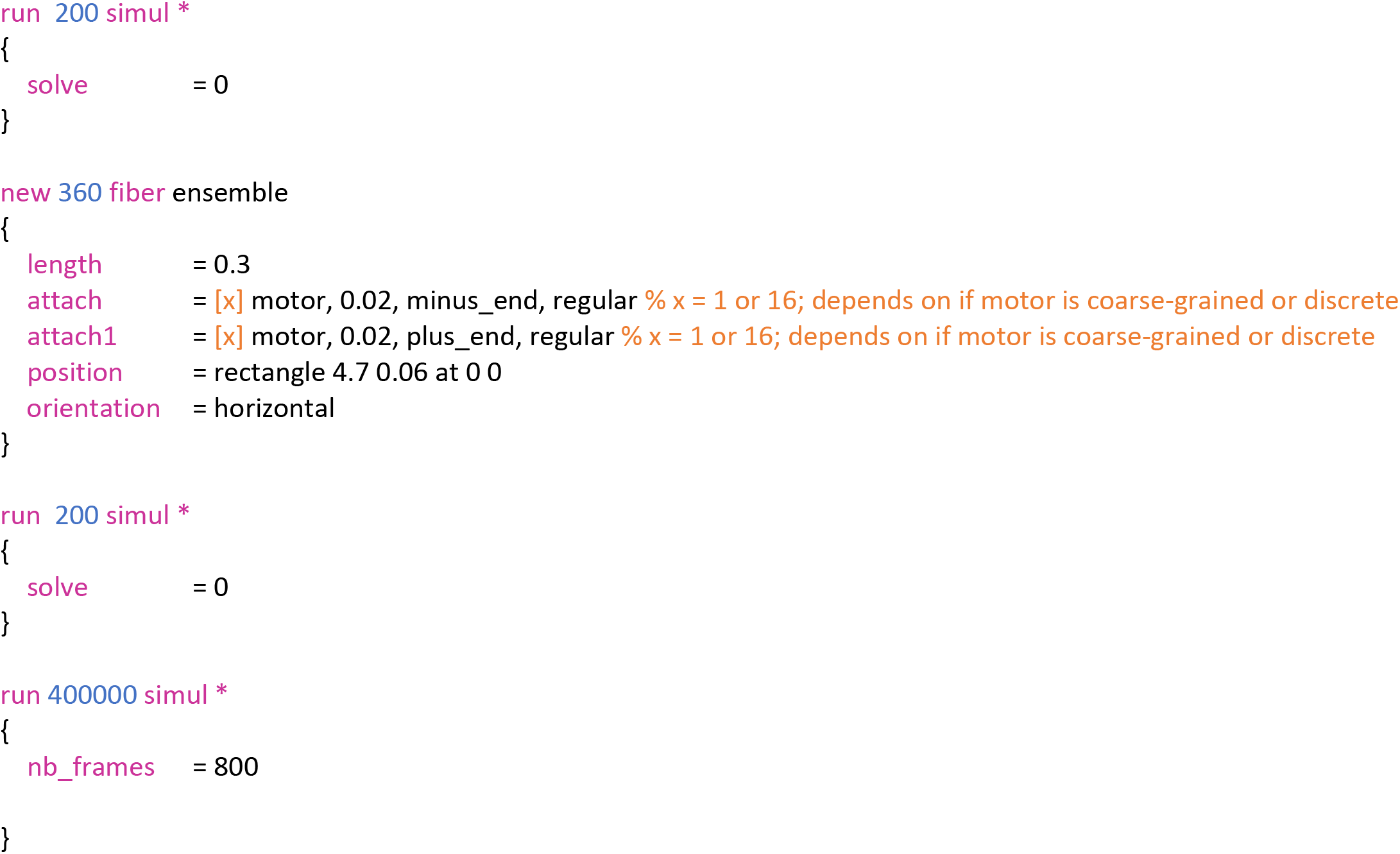

Parameters of motors are shown for each bond-type and for discrete vs coarse-grained versions of all motor ensembles.

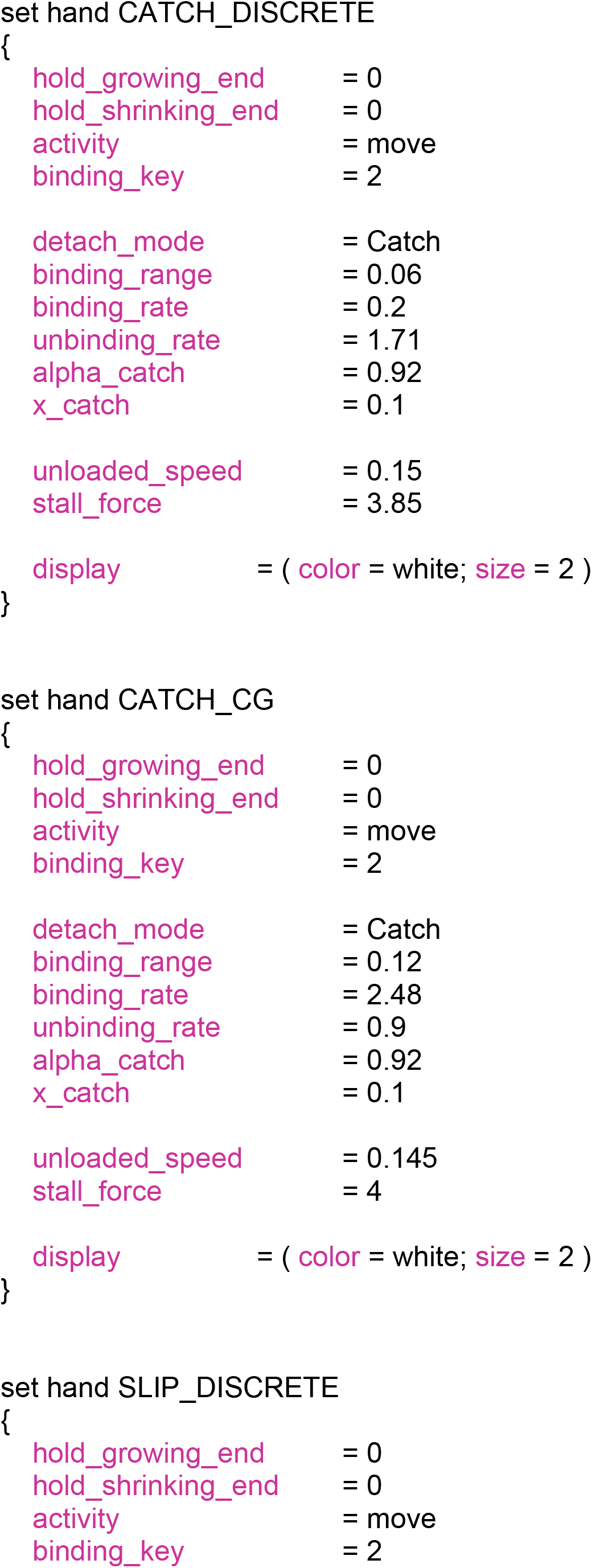
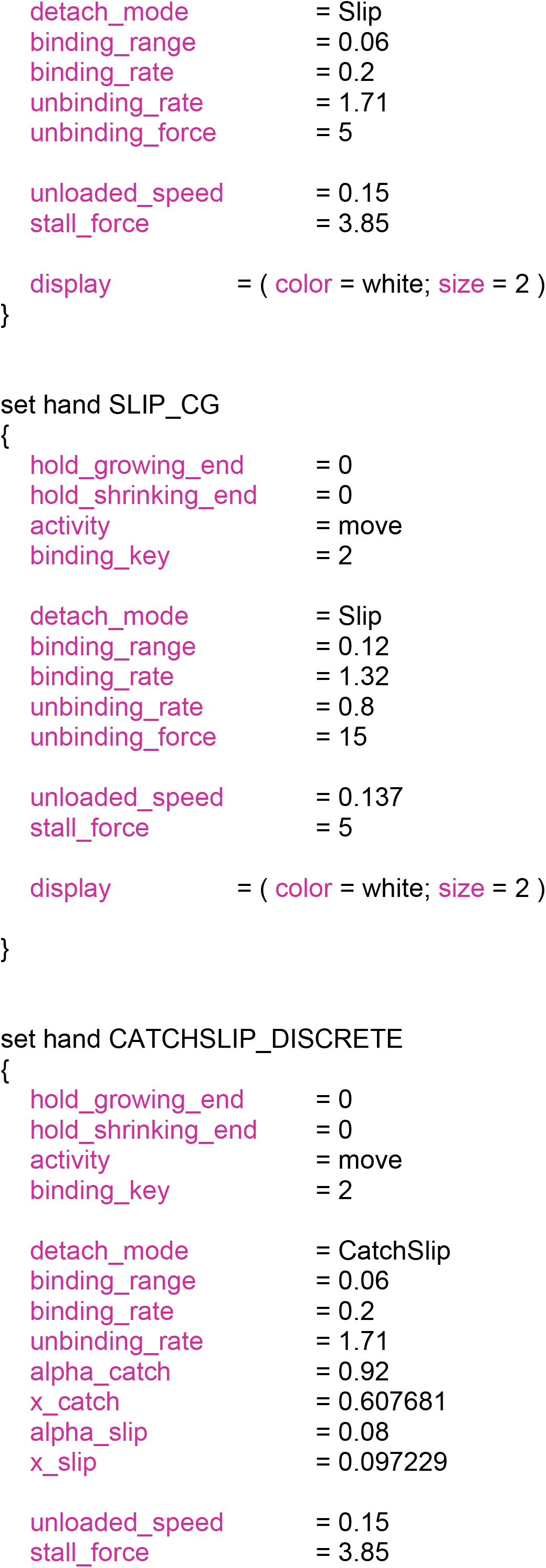
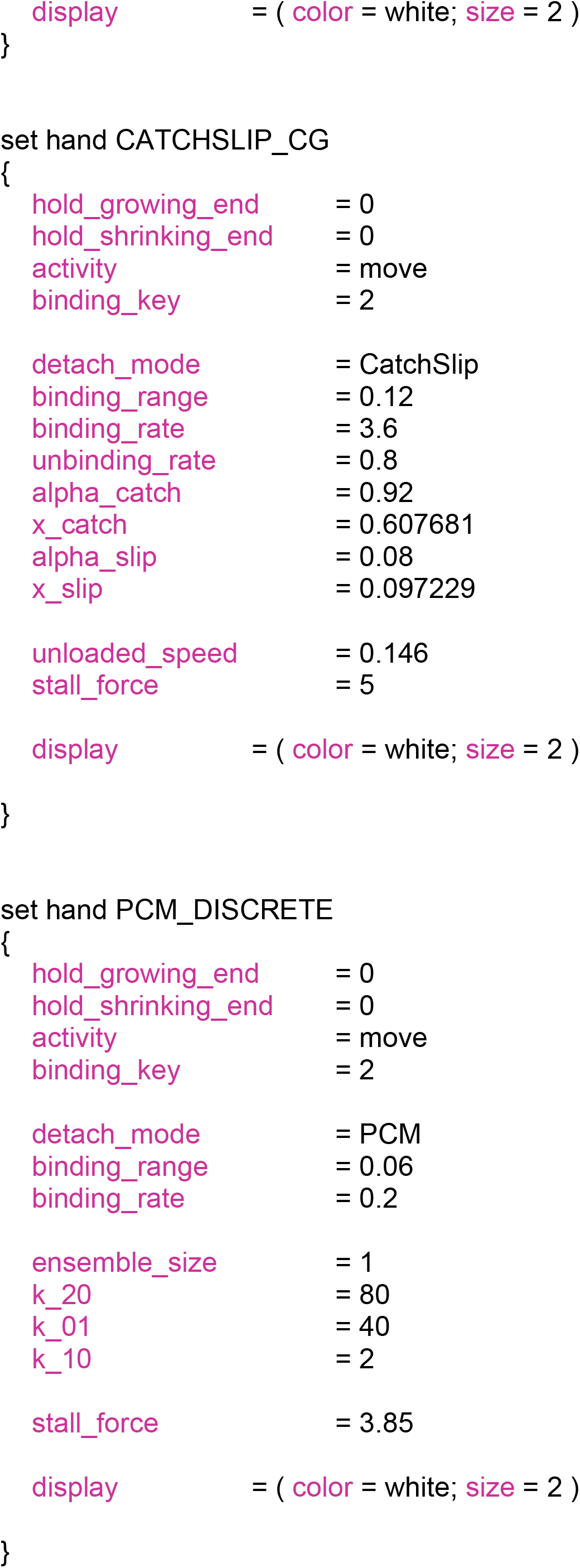
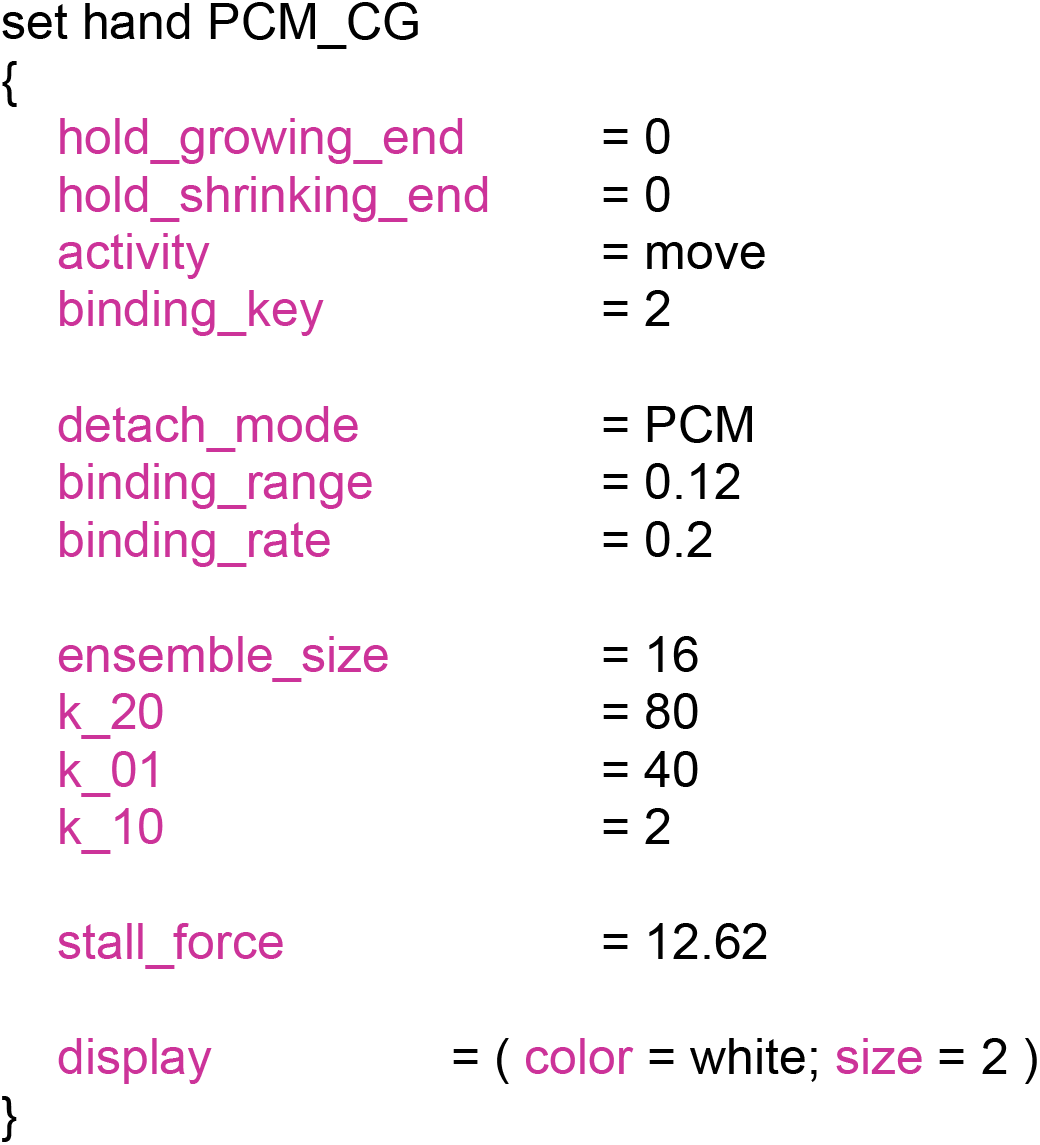

